# Lipid fingerprints are similar between SLC6 transporters in the neuronal membrane

**DOI:** 10.1101/2021.01.20.427530

**Authors:** Katie A. Wilson, Lily Wang, Yie Chang Lin, Megan L. O’Mara

## Abstract

We use molecular dynamics simulations to characterise the local lipid annulus, or “fingerprint”, of three SLC6 transporters (dDAT, hSERT, and GlyT2) embedded into a complex neuronal membrane. New membrane analysis tools were created to improve leaflet detection and leaflet-dependent properties. Overall, lipid fingerprints are comprised of similar lipids when grouped by headgroup or tail saturation. The enrichment and depletion of specific lipids, including sites of cholesterol contacts, varies between transporters. The subtle differences in lipid fingerprints results in varying membrane biophysical properties near the transporter. Through comparisons to previous literature, we highlight that the lipid-fingerprint in complex membranes is highly dependent on membrane composition. Furthermore, through embedding these transporters in a simplified model membrane, we show that the simplified membrane is not able to capture the biophysical properties of the complex membrane. Our results further characterise how the presence and identity of membrane proteins affects the complex interplay of lipid-protein interactions, including the local lipid environment and membrane biophysical properties.

**HIGHLIGHTS:**

- Lipid fingerprints are comprised of similar lipid classes
- Sites of specific lipid contacts, including CHOL, varies between transporters
- Changes in lipid annulus result in variable local membrane biophysical properties
- Membrane composition, including that of complex membranes, affects lipid annulus

**GRAPHICAL ABSTRACT:** 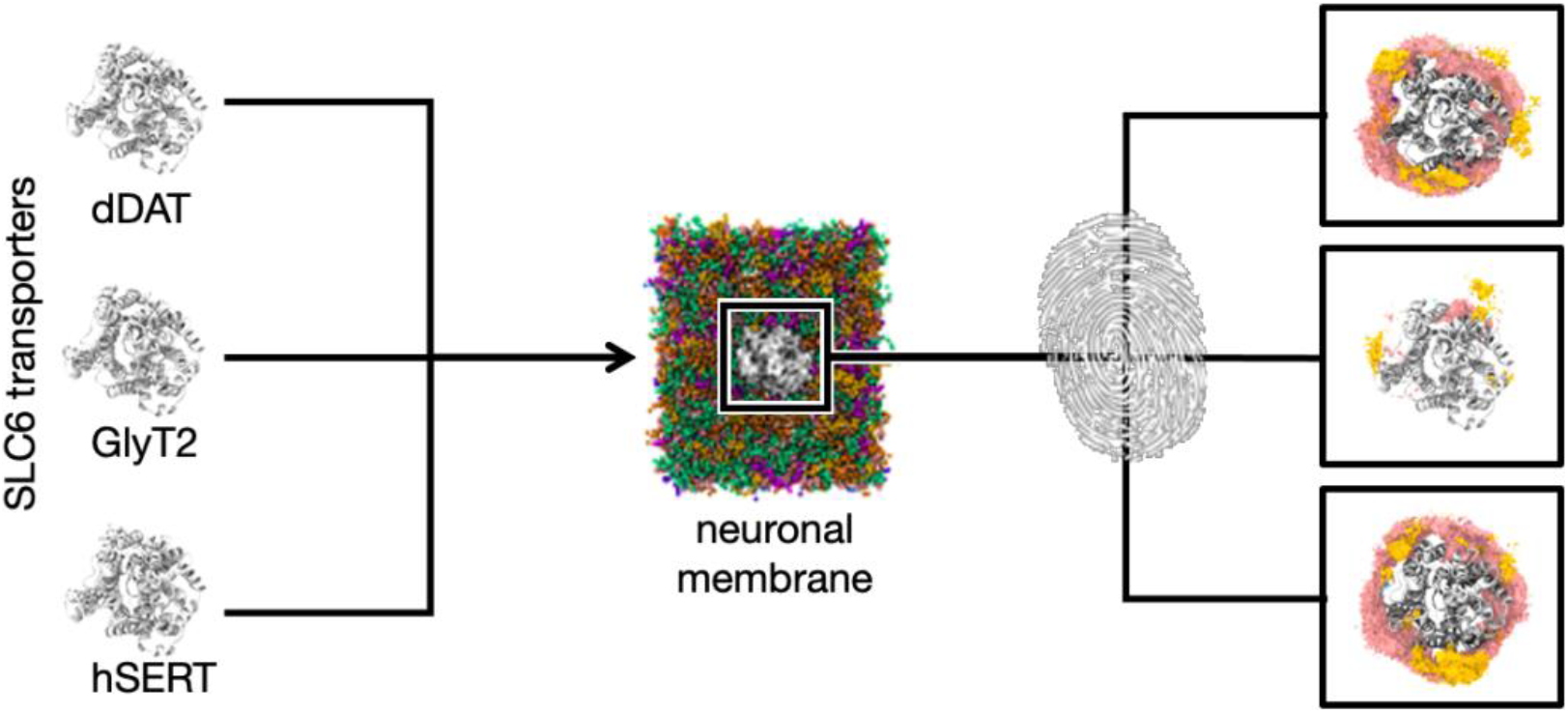

## INTRODUCTION

The composition of cell membranes encompasses tremendous chemical diversity. A typical bilayer contains hundreds of different lipid species, distributed asymmetrically between the two leaflets [1]. Membrane environments are also constantly changing, as lipids mix laterally to form distinct domains and translocate between leaflets in a variety of biological processes [2]. The complex and dynamic lipid environment gives rise to correspondingly complex interactions with embedded membrane proteins. Lipids play roles in protein trafficking, localization, and activity [3]. They modulate membrane protein behaviour both specifically, by binding to the protein at distinct binding sites [4], and non-specifically, through changes in the biophysical properties of the surrounding membrane environment that in turn affects the protein function [5]. Several properties are known to affect protein behaviour, including the thickness of the membrane, the lateral pressure field of the membrane, and the organization of charges at the protein-lipid interface [6].

A typical membrane lipid is comprised of a polar headgroup and one or more acyl chain tail groups, and the combinatorial possibilities result in membranes that incorporate extensive chemical diversity [1]. Headgroups common to mammalian membranes include phosphatidylcholines (PC), phosphatidylethanolamines (PE), phosphatidylserines (PS), phosphatidylinositols (PI), and phosphatidic acid (PA). In addition, PI lipids may be phosphorylated to form phosphorylated PIs (PIPs), while PC and PE headgroups can be combined with a sphingosine backbone to form sphingolipids (SM). Alternatively, the headgroup can be a hydroxyl group, creating ceramides (CER), or a carbohydrate, creating glycosphingolipids (GS). GS lipids are typically found only in the extracellular leaflet, and PI, PA and PS lipids in the intracellular leaflet. The tails join phosphoglycerides at two positions, denoted *sn-1* and *sn-2*. Acyl chains range in length from 16–22 carbons in synaptic membrane phospholipids [7], and vary in saturation from saturated to up to 6 *cis* double bonds, with further variation in the position of these double bonds. The tails join phosphoglycerides at two positions, denoted *sn-1* and *sn-2*. In mammals, the *sn-1* tail is commonly unsaturated, whereas the *sn-2* tail is often mono- or polyunsaturated [8]. Sterols are also present in membranes, most commonly as cholesterol (CHOL) [1].

The combinatorial complexity of membranes can be difficult to study at a molecular level using experimental techniques, which typically yield ensemble-averaged data. Molecular dynamics (MD) simulations probe membrane and transmembrane protein interactions at a molecular resolution, enabling us to study time-dependent behaviour and quantify subtle changes to membrane properties. Coarse-graining, the practice of representing multiple atoms with one particle “bead”, allows the simulation of larger systems for longer timescales than atomistic simulations. The balance between fine spatiotemporal resolution and breadth of accessible size and timescales has made coarse-grain (CG) MD a popular technique for quantitative studies of membrane behaviour [9].

Recent studies have used CG MD to model an average plasma membrane [10] and a neuronal membrane [11] from experimentally determined lipid compositions. These are the first computational models to approach the complexity characteristic of realistic membranes. Until the publication of these models, MD simulations of transmembrane proteins incorporated at most four or five different lipid species [9]; the neuronal and plasma membrane models each contain over fifty lipid species. The average properties of the neuronal membrane model are comparable with those of the plasma membrane in properties such as bilayer thickness, tail order, diffusion, and flip-flop, although diffusion and flip-flop rates are slightly slower in the neuronal model. However, changes in properties in the local environment are likely more meaningful for lipid-protein interactions than the average property of the membrane. A previous study examining ten different classes of membrane proteins embedded in the average plasma membrane showed that each protein induced changes in the local membrane environment to form a lipid annulus unique to that protein class. The distinct, non-uniform distributions of different lipid types around each protein perturb local membrane properties to form a protein-specific “lipid fingerprint” [12]. These local membrane properties are highly dependent on lipid composition. Another study compared the complex neuronal membrane to a series of model membranes of decreasing chemical complexity; the results suggest that a complex model incorporating diverse lipid species is important for the accurate representation of a number of properties that can modulate transmembrane protein behaviour, such as membrane fluidity, cholesterol localization, lipid clustering, and membrane curvature [13]. Therefore, the lipid fingerprints observed around each protein in an averaged membrane may not be representative of the true environment that may emerge in a tissue-specific membrane.

One of the proteins examined in the previous plasma membrane study was the dopamine transporter (DAT) [12], a neurotransmitter transporter from the solute carrier 6 (SLC6) family. SLC6 neurotransmitter transporters are responsible for neurotransmitter reuptake from the synaptic cleft. Dysregulation of SLC6 neurotransmitter transporters leads to disruption of neurotransmission and has been implicated in a range of disorders including Parkinson’s disease, addiction, depression, chronic pain, and epilepsy [14]. Here we examine the lipid environment obtained when DAT and two other SLC6 neurotransmitter transporters, hSERT (48.61% sequence identity to dDAT) and GlyT2 (45.76% sequence identity to dDAT), are embedded in a complex neuronal membrane. We investigate whether lipid fingerprints are unique to members within the same protein family; whether the local lipid environment is substantially different in the neuronal membrane compared to the plasma membrane; and the effect of the presence of a protein in the neuronal membrane on membrane properties. We then embed the same proteins in a simplified model membrane (POPC/CHOL) comprised of 80% 1-palmitoyl-2-oleoyl-sn-glycero-3-phosphocholine (POPC) and 20 % cholesterol, to explore whether the presence of membrane protein mitigates or modifies the known changes in bilayer properties between simple and complex membranes.

## METHODS

All simulations were prepared and performed using the 2016.1 version of the Groningen Machine for Chemical Simulation (GROMACS) [15] and version 2.2 of the Martini forcefield [16]. The coordinates of the *Drosophila melanogaster* dopamine transporter, dDAT (PDB ID: 4XP1) [17], and human serotonin transporter, hSERT (PDB ID: 5I6X) [18], in the outward-occluded conformation were obtained from the Protein Data Bank. The homology model of GlyT2 in the outward-occluded conformation was obtained from Subramanian *et al*. [19]. Mutations to the wild-type sequence that were introduced to help with crystallography were reversed (specifically, V50A and L350A in dDAT; and I218A, T366S, C418A and C507A in hSERT). GlyT2, dDAT and hSERT were coarse-grained using the *martinize* protocol [20]. Each coarse-grained protein was then inserted into a lipid bilayer generated using a custom version of the *insane* script [21]. The model two-component bilayer comprised 80% phosphatidylcholine (POPC) and 20% cholesterol while the neuronal membrane was set up using previously reported lipid compositions (Table S1) [22]. Parameters for the lipids were obtained from the same paper [22]. The system was solvated using polarizable coarse-grained water. Each system contained ~930 lipids in a box size of 16.6 × 19.4 × 16 nm with the membrane oriented in x-y plane. NaCl was added to a physiological concentration of 0.15 M. Additional ions were added to give overall charge neutrality for each system.

All 6 systems were energy minimized using a steepest descent algorithm. The systems were then equilibrated by performing a series of five sequential 1 ns simulations with stepwise 1000, 500, 100, 50 or 10 kJ mol^−1^ nm^−2^ position restraints on the protein. Each system was simulated in triplicate for 10 μs, using a 20 fs timestep, in the recommended configuration for MARTINI [23]. Random velocities were assigned at the beginning of each simulation to initiate the simulation. The lengths of the covalent solute bonds were constrained using the LINCS algorithm. Simulations were performed in the NPT ensemble, with the temperature maintained at 310 K using the Bussi-Donadio-Parrinello velocity-rescaling thermostat [24] with a coupling constant of τ_t_ = 1 ps. The pressure was maintained at 1 bar through coupling to a Parrinello-Rahman barostat [25] with a coupling constant of τ_p_ = 12 ps and an isothermal compressibility of 3 × 10^−4^ bar. The non-covalent interactions were calculated with a 1.1 nm cut-off.

Analysis was performed on frames at 10 ns intervals. The membrane thickness was calculated with *g*_*thickness* [26], using the PO4 and GM1 beads of lipids. Lipid diffusion in the xy plane was calculated using *gmx msd* on the final 5 μs of the simulation. The 200 to 500 ns lag-time portion of the MSD was used to avoid ballistic trajectories obtained at short lag-times and poor averaging obtained at long lag-times [27]. To characterize how lipids within the complex neuronal membrane bind to the neurotransmitter transporters, we calculated residues within 6 Å of a lipids using the Visual Molecular Dynamics (VMD) software [28]. Lipid contact fractions were calculated as described in Wilson *et al., 2020* [13]. All other membrane analyses (leaflet detection, flip-flop rates, area per lipid, and depletion-enrichment index calculation) were carried out using Python code built upon the *MDAnalysis* package [29,30], available at https://github.com/OMaraLab/SLC6_lipid_fingerprints (details in SI).

## RESULTS AND DISCUSSION

### The local lipid environment is similar around different SLC6 membrane proteins

We characterize the lipid annulus around each SLC6 transporter with lipid depletion-enrichment and contact analyses. Lipid depletion-enrichment indices (DEI) summarize the enrichment or depletion of a certain lipid group within the lipid annulus around the protein, compared to the composition of the bulk membrane, over the simulation. We use a lipid annulus of 6 Å from the edge of the SLC6 transporter, with a 2 Å buffer around this annulus, to quantify the immediate lipid environment. To identify specific lipid-protein interactions, we examine which lipid groups form contacts, defined as being within a distance of ≤ 6 Å, with specific regions of each transporter that persist for >40% of each trajectory. In general, the local lipid environment around each SLC6 transporter is similar across major lipid headgroup types, although differences arise when inspecting the interactions between the transporter and individual lipid species. We explore the behaviour of lipids both by headgroup and tail saturation.

#### The lipid annulus around dDAT, GlyT2, and hSERT is enriched in polyunsaturated lipids

The neuronal membrane contains saturated, monounsaturated, and polyunsaturated species (Figure 1 and Table S1). These are distributed asymmetrically across the intracellular and extracellular leaflet. On average, lipids in the extracellular leaflet contain 1.76 unsaturated bonds, while the intracellular leaflet contains an average of 3.25 unsaturations per lipid. Previous studies have shown that endogenous polyunsaturated fatty acids and their synthetic derivatives regulate the activity of SLC6 transporters, in particular GlyT2 [31] and the dopamine transporter [32,33]. Polyunsaturated lipids are enriched around GlyT2 and dDAT in both leaflets (DEI_extracellular_ = 1.284 ± 0.184, DEI_intracellular_ = 1.257 ± 0.061, Figure 2), although only to a minor extent around hSERT in the extracellular leaflet. For each embedded SLC6 transporter, saturated lipids are depleted both around the protein (DEI_extracellular_ = 0.475 ± 0.093; DEI_intracellular_ = 0.432 ± 0.079), and in the vicinity of polyunsaturated lipids and cholesterol (Figure 3). Monounsaturated lipids are not significantly enriched nor depleted in the extracellular leaflet; they are slightly depleted in the intracellular leaflet (DEI_intracellular_ = 0.715 ± 0.149).

**Figure 1.**
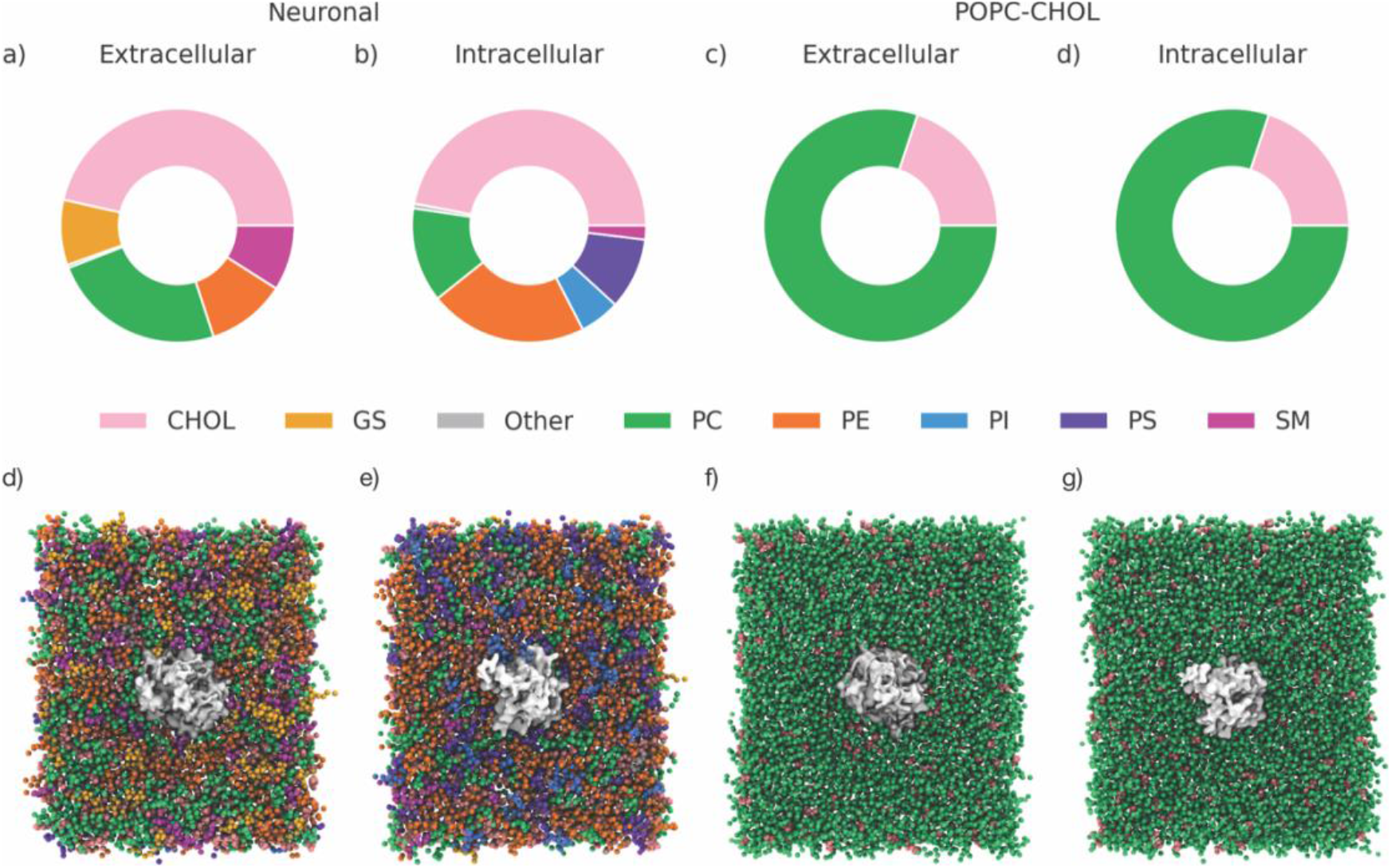
Membrane composition by headgroup (top: a–d) and starting orientation of protein-bilayer systems (bottom: d–g) for neuronal system (left: a, b, d, e) and the model POPC-CHOL system (right: c, d, f, g). Proteins are shown in grey. Full composition and definition of acronyms are given in Table S1.

**Figure 2.**
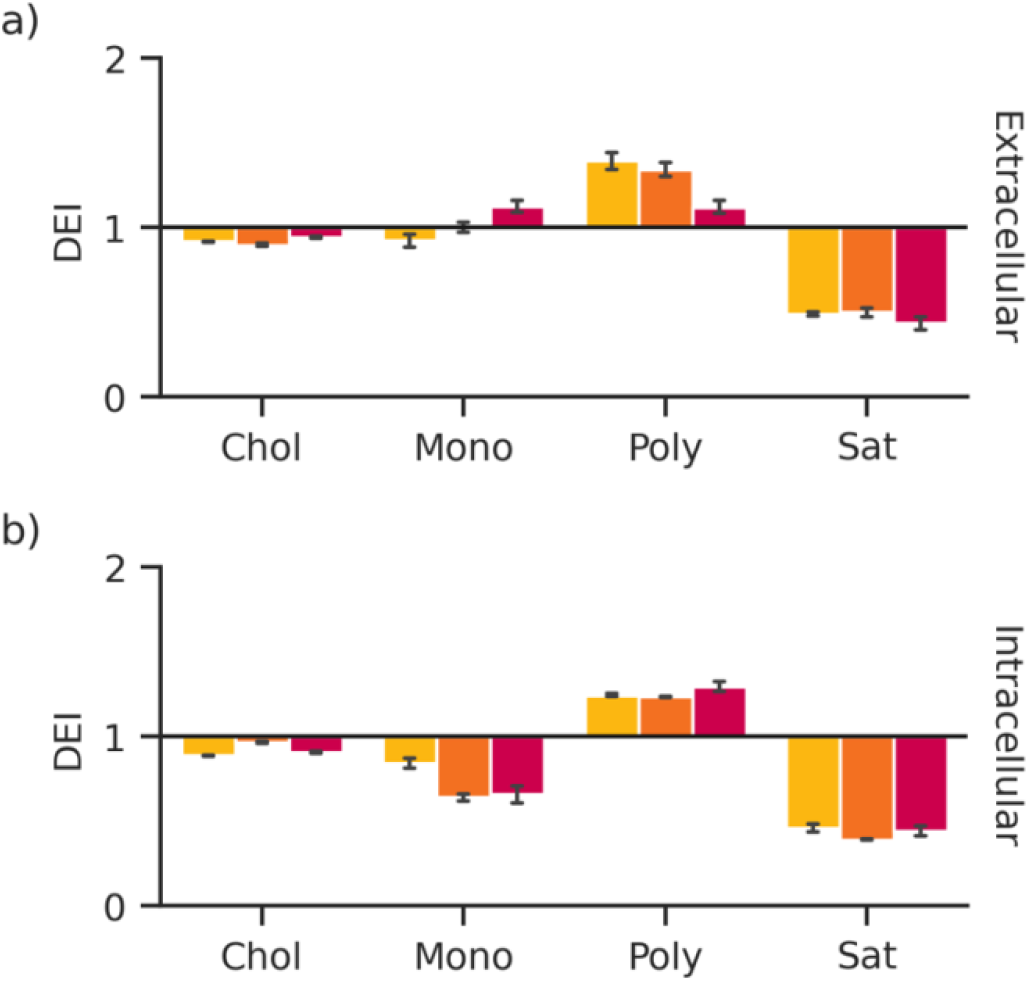
The mean depletion-enrichment index (DEI) of lipids in the protein annulus grouped by tail saturation, in the **a)** extracellular leaflet; and **b)** the intracellular leaflet. The standard error from the mean is represented by the error bars. DEI values of each lipid species are shown in Table S2. Yellow: dDAT. Orange: GlyT2. Red: hSERT.

**Figure 3.**
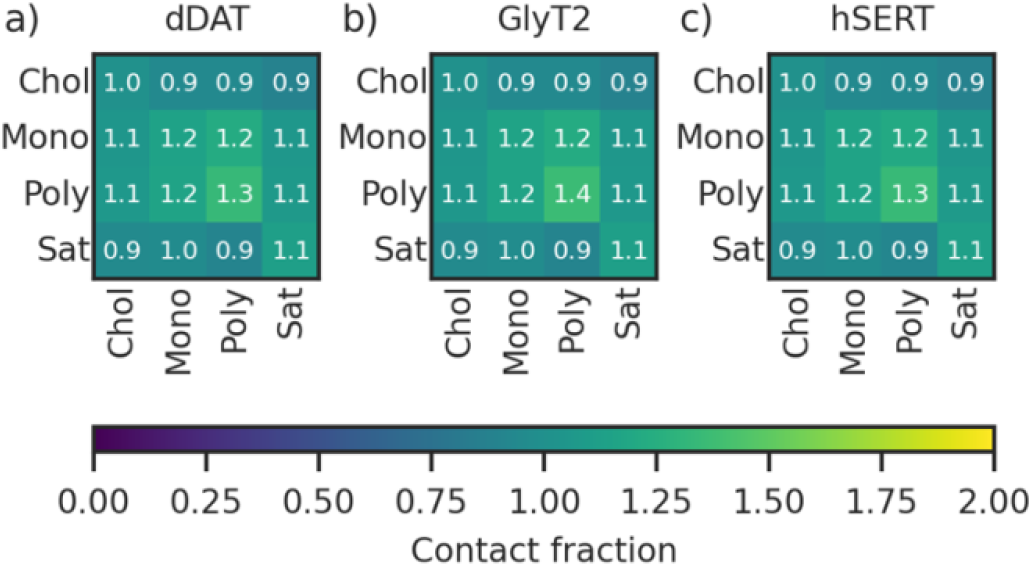
The average lipid contact fractions of the neuronal membrane, by tail saturation, for membranes with embedded: **a)** dDAT, **b)** GlyT2, and **c)** hSERT.

#### Phosphatidylcholine and phosphatidylserine lipids form few and non-specific sustained contacts with SLC6 transporters

Phosphatidylcholines (PC) are the most abundant class of glycerophospholipid in the neuronal membrane (24% of the extracellular and 13% of the intracellular leaflet), and the second most abundant group of lipids after cholesterol. Six polyunsaturated species (PAPC, DOPC, PUPC, PFPC, OIPC, OUPC), one monounsaturated species (POPC) and one saturated species (DPPC) are present in the neuronal membrane. Despite their abundance in the membrane, as a class, PC lipids are largely depleted in the lipid annulus around the SLC6 transporters in both the intracellular and extracellular leaflets of the membrane (DEI_extracellular_ = 0.781 ± 0.091; DEI_intracellular_ = 0.630 ± 0.073, Figure 4). The population of PC lipids within the membrane appears to influence which particular lipid species form significant contacts with the transporter, as persistent contacts are only formed with lipids that constitute at least 1% of the membrane. These contacts are formed with different regions of each transporter, indicating no overall pattern or specificity in the contacts formed. For example, a single POPC lipid forms direct contacts with dDAT for up to 19% of the total simulation time (30 μs) in the region of TM5, while PUPC and PAPC lipids form transient interactions with TM8 that collectively account for ~30% of the total simulation time. Collectively, PAPC lipids interact with GlyT2 in the region of EL3 and TM11 or TM12, across 2 of the 3 replicate simulations. These interactions each persist for up to 59% of a single 10 μs replicate simulation. Similarly, in the case of hSERT, POPC lipids collectively interact with TM7, while PAPC lipids collectively interact with the hSERT N-terminus, each for 40% of a single 10 μs replicate simulation. However, no single PC lipid forms significant contacts (>40 % simulation time) with either GlyT2 or hSERT.

**Figure 4.**
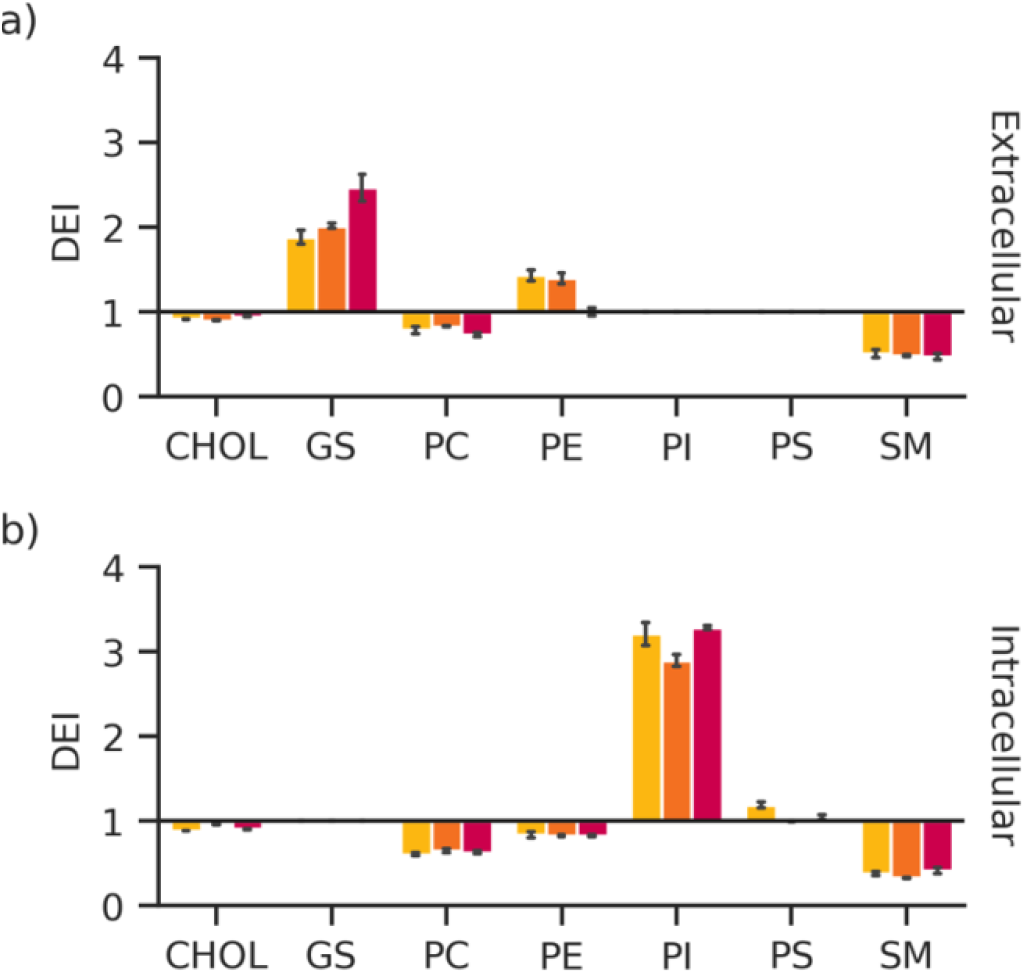
The mean depletion-enrichment index (DEI) of lipids in the protein annulus grouped by head group, in the **a)** extracellular leaflet; and **b)** the intracellular leaflet. The standard error from the mean is represented by the error bars. DEI values of each lipid species are shown in Table S2. Yellow: dDAT. Orange: GlyT2. Red: hSERT.

Phosphatidylserine (PS) lipids are only found in the intracellular leaflet and comprise 10% of all lipids in the intracellular leaflet. As a class, PS lipids are neither enriched nor depleted around each transporter (DEI_intracellular_ = 1.079 ± 0.119). However, this analysis averages the enrichment behaviour of each PS lipid (Figure S2). Two highly unsaturated lipid species are enriched around each transporter: OUPS (DEI = 1.311 ± 0.515) which contains 7 unsaturations (C18:1/22:6 tails), and PUPS (DEI = 1.344 ± 0.247) containing 6 unsaturations (C16:0/22:6). Notably, the latter lipid does not form many sustained contacts despite its enrichment and only comes into contact with each of the SLC6 transporters for 40–44% of a single 10 μs replicate simulation per protein. In contrast to OUPS and PUPS, the saturated DPPS (DEI = 0.447 ± 0.352) and monounsaturated POPS (DEI = 0.667 ± 0.186) are generally depleted around each transporter. PAPS, a polyunsaturated lipid with 4 unsaturations (C16:0/20:4) is enriched around both dDAT (DEI = 1.300 ± 0.263) and hSERT (DEI = 1.265 ± 0.193), but neither enriched nor depleted around GlyT2 (DEI = 0.962 ± 0.163). Persistent contacts (lasting for >40% of any single trajectory) are formed with residues in TM3, IL4, TM9, and TM12 of dDAT, but none are formed with hSERT or GlyT2. Overall, PS lipid headgroups are most enriched around dDAT, while DPPS, PAPS, POPS, and PUPS are most depleted around hSERT.

#### Phosphatidylethanolamines interact differentially with dDAT, GlyT2, and hSERT through π-interactions with the lipid tail

Phosphatidylethanolamine (PE) lipids are the second most abundant glycerophospholipid in the neuronal plasma membrane. The neuronal membrane contains five polyunsaturated species (PUPE, PAPE, OAPE, OUPE and OIPE) and a single monounsaturated (POPE) species of PE lipids. PE lipids are the only lipid family to have different depletion-enrichment profiles over the intracellular and extracellular leaflets. PE lipids comprise 22% of the intracellular leaflet. Despite their relative abundance in the intracellular leaflet, PE lipids are depleted around all three SLC6 transporters in this leaflet (DEI_intracellular_ = 0.831 ± 0.077). The enrichment varies by protein in the extracellular leaflet, where PE lipids comprise 11 mol % of the total lipids; the relative depletion/enrichment index shows that they are slightly enriched around dDAT (DEI_extracellular_ = 1.434 ± 0.243) and GlyT2 (DEI_extracellular_ = 1.397 ± 0.245), but not hSERT (DEI_extracellular_ = 1.004 ± 0.165). The lipid enrichment profiles of individual PE lipid species also vary by both protein and leaflet (Figure S2). For example, lipids in the extracellular leaflet such as OAPE, PUPE and OAPE are the most enriched around dDAT and the least around hSERT, following the trend of the overall PE class. Inversely, lipids in the intracellular leaflet such as OAPE and OIPE are the most depleted around dDAT but enriched around hSERT.

The contact profiles of lipid-protein interactions correlate with the DEI trends in the extracellular leaflet, but highlight differences in lipid interaction behaviour between the three SLC6 transporters (Figure S3). We defined any interaction between each protein residue and a particular lipid species to be a “persistent” contact if it occurred >40% any single trajectory. dDAT forms the greatest number (112 contacts) of persistent contacts with PE lipids, followed by GlyT2 (71 contacts). hSERT forms the fewest persistent contacts (30 contacts). Furthermore, the contacts are maintained for longer in each 10 μs replicate for dDAT (maximum 80% and median of 51% of each 10 μs simulation), than for GlyT2 (max 68% and median 47%) or hSERT (max 60% and median 44%). Most of the interactions with PE lipids take place in helical regions of each transporter (dDAT: 79%, GlyT2: 75% and hSERT: 60%) and do not occur as frequently in loop regions. The most frequently occurring contact sites of PE lipids for dDAT are in TM3 (12%), TM5 (12%), TM10 (13%) and TM12 (17%) (Figure 5a and 6); in GlyT2, 27% of the PE contacts occur on TM11 and an additional 13% of the contacts occur on each TM5 and EL3. PE contacts with hSERT most frequently occur with IL3 (17%), and TM7 (17%). While there is some overlap in the most prevalent sites of PE contacts for each of the investigated SLC6 transporters, there are also significant differences between these sites of interaction. This suggests that although lipid fingerprints of proteins within the same family may be broadly similar, unique differences may emerge at the level of interaction with individual lipid species.

**Figure 5.**
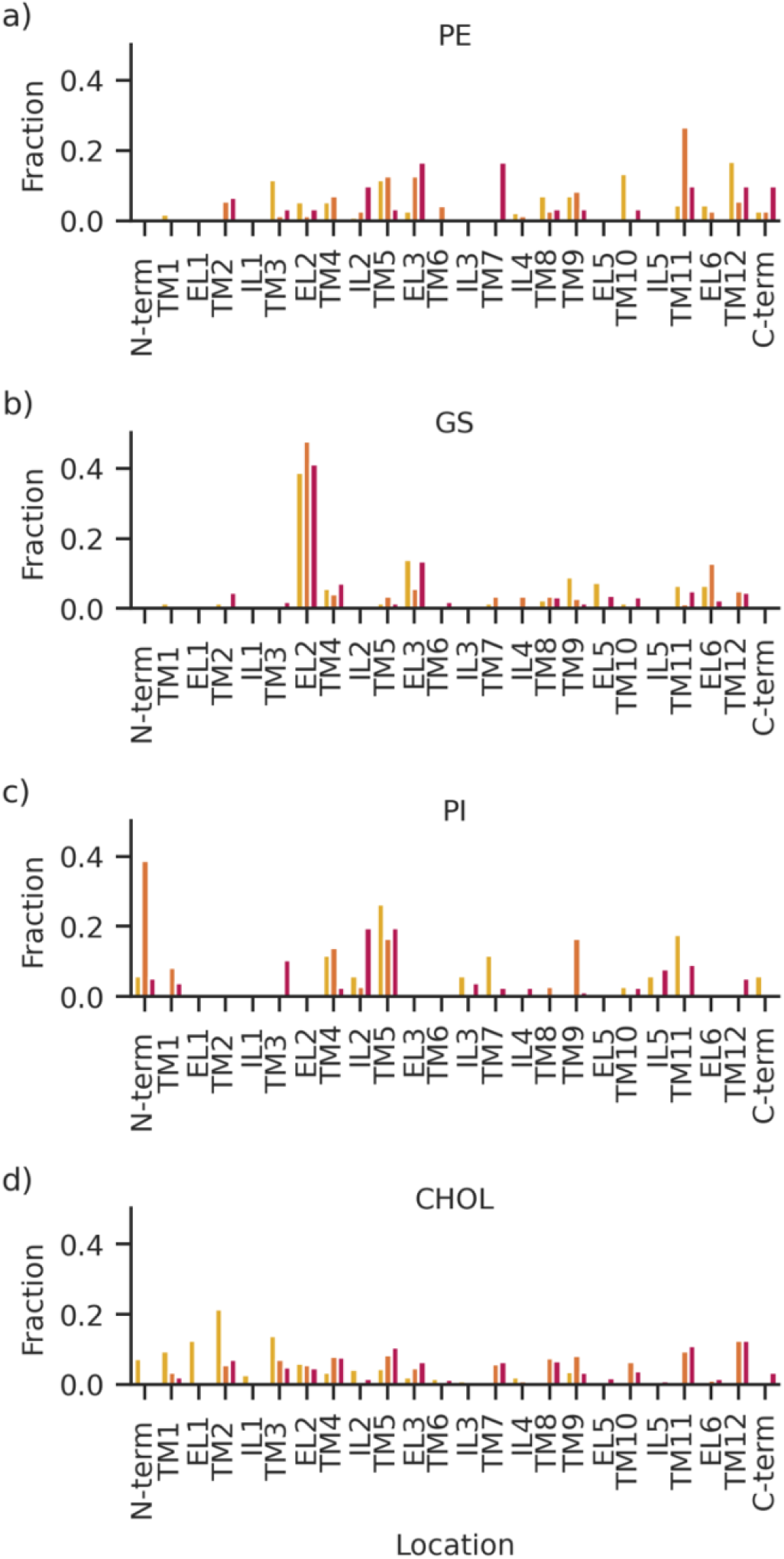
The fraction of contacts formed by each class of lipids with the protein, by protein region. **a)** PE lipids; **b)** GS lipids; **c)** PI lipids; **d)** cholesterol. Yellow: dDAT. Orange: GlyT2. Red: hSERT.

**Figure 6.**
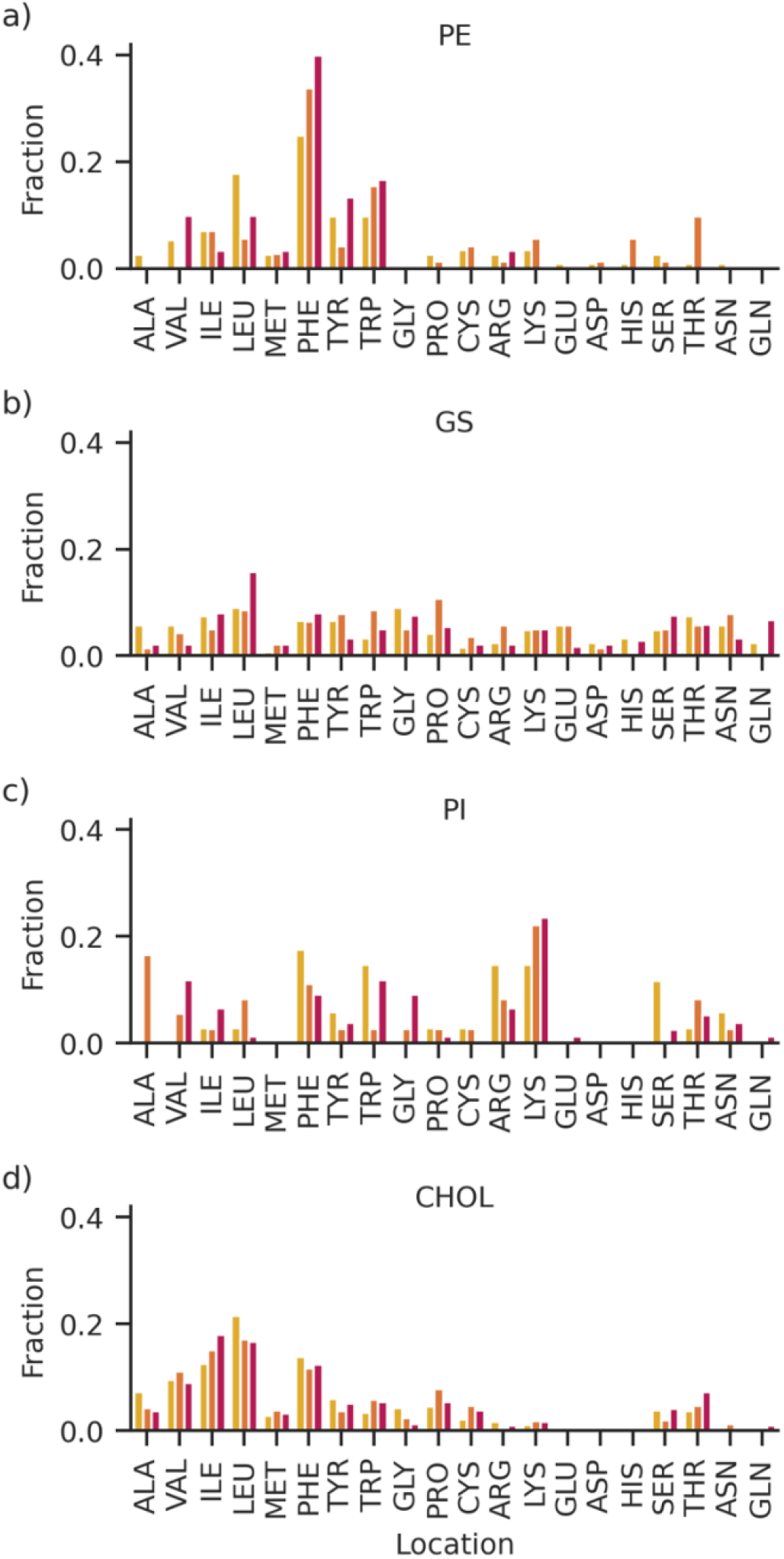
The fraction of contacts formed by each class of lipids with the protein, by amino acid. **a)** PE lipids; **b)** GS lipids; **c)** PI lipids; **d)** cholesterol. Yellow: dDAT. Orange: GlyT2. Red: hSERT.

PE lipids also interact preferentially with certain amino acids, as shown in Figure 6a. Across all three SLC6 transporters, protein-PE lipid contacts preferentially form with phenylalanine (40%, 34% and 28% of all contacts for hSERT, GlyT and dDAT respectively). Phenylalanine comprises between 7.9% (hSERT) and 9.0% (dDAT) of the total amino acids in each of the three transporters (Figure S4). PE lipids also form frequent contacts with tryptophan in both dDAT (10% of contacts) and hSERT (17% of contacts), despite tryptophan comprising only 3.2–3.5% of the amino acids of these transporters. PE lipids interact with glycine and alanine to a lesser extent than may be expected from their relative abundance within each transporter. The prevalence of tail group interactions with aromatic groups suggests that the double bonds within the highly unsaturated lipid tails are forming CH-π or π-π interactions with the aromatic amino acids. These non-covalent interactions have been noted to provide stability of up to ~30 kJ/mol to protein-protein [34], nucleic acid-protein [35,36] and carbohydrate-protein complexes [37,38] but have not been previously identified in lipid-protein interactions [39].

Overall, the PE contacts with the transporter are very dynamic and PE lipids exchange rapidly in each binding site. dDAT is the only transporter where individual PE species bind to the transporter for >40% of the 30 μs of total simulation time, but the binding site of these PE lipids varies between replicates. The PE contacts are driven almost entirely by tail group interactions; only 7–8% of the contacts with dDAT and GlyT2 occur with a headgroup, and none of the PE contacts with hSERT are with the headgroup. The preference of these protein-tail lipid interactions towards PE lipids is striking as the tail groups are present with other headgroups as well (e.g., PUPC, PAPC, OUPC, PUPS and OUPS) but these lipids do not form persistent contacts with the transporters. The high occupancy of PE lipids around the SLC6 transporters may occur due to the negative curvature induced by the small lipid headgroup and polyunsaturated tails, which can curve the membrane to suit the hydrophobic depth of the transporter.

#### Glycosphingolipids form long-lived headgroup interactions with dDAT, GlyT2, and hSERT loop regions

Glycosphingolipids (GS) make up 9% the extracellular leaflet of the model neuronal membrane and include glucosylceramides and glycosylated gangliosides. Glycosphingolipid species in the neuronal membrane comprise the monounsaturated (DPGS and DBGS) and polyunsaturated glucosylceramides (PNGS and POGS), monounsaturated monosialotetrahexosylgangliosides (DPG1) and the monounsaturated monosialodihexosylgangliosides (DPG3). As a group, glycosphingolipids are enriched around each transporter but more so around hSERT (DEI = 2.466 ± 0.579; 235 persistent contacts) than in comparison to dDAT (1.882 ± 0.296; 121 persistent contacts) or GlyT2 (2.013 ± 0.105; 140 persistent contacts). Individually, the DEI and lipid contact profiles vary between transporters. The enrichment profiles of individual lipid species vary around each transporter and often between replicates simulations for the same transporter, as extreme DEI values are driven by the low population of the glycosphingolipids. The most common glycosphingolipid, DPGS, comprises 4.9% the extracellular leaflet. It is consistently enriched around every transporter, across all replicate simulations. Greater enrichment of DPGS is observed around hSERT (DEI = 2.641 ± 0.639) than dDAT (DEI = 1.727 ± 0.398) and GlyT2 (DEI = 1.761 ± 0.451) (Figure 2). Lower population glycosphingolipids vary much more; they are not systematically enriched or depleted around any of the three SLC6 transporters but are both enriched and depleted in different replicates of at least one transporter. PNGS forms 1.1% the extracellular leaflet and is enriched around hSERT in all three replicates (DEI = 2.043 ± 0.282), but its interaction with dDAT and GlyT2 is variable: the low concentration (and consequently, low absolute number of molecules) of PNGS within the membrane results in it being both significantly enriched and depleted around dDAT (DEI = 2.272 ± 1.965) and GlyT2 (DEI = 1.404 ± 1.580) in different replicates. Four molecules each of DPG1, DPG3, and DBGS are in the membrane, constituting 0.9% the extracellular leaflet, while two molecules of POGS comprise 0.4% the extracellular leaflet. The aggregation of these lipids around dDAT is also heavily dependent on each individual simulation replicate: they are severely depleted in at least one replicate but enriched in another (Table S2). The same variance is observed to a lesser extent with hSERT, where DBGS, DPG3, and POGS are all enriched or depleted in different replicates (Table S2). In contrast, DPG1 (DEI = 4.812 ± 2.634) is enriched around hSERT across all three replicates. In the case of GlyT2, the aggregation of GS lipids is more systematic: DBGS is depleted in each 10 μs replicate simulation (DEI = 0.656 ± 0.384), while DPG1, DPG3 are each enriched in each replicate simulation (DEI_DPG1_ = 3.254 ± 1.376, DEI_DPG3_ = 4.523 ± 2.487, DEI_POGS_ = 1.644 ± 0.910). The variation in individual lipid profile underscores that lipid annulus is a dynamic environment and that differences in the average lipid fingerprint at the individual lipid species level (particularly for low abundancy species) may arise from the stochastic nature of lipid diffusion.

The contacts formed by lipids to each transporter suggest that the nature of interactions differs between each of the GS lipids (Figure S5). Individual glycosphingolipids containing polysaccharide headgroups (G1 and G3) form more long-lived contacts (≥ 70% in any single trajectory) than individual lipids with monosaccharide headgroups. DPG1 forms long-lived contacts with two replicates of dDAT and all replicates of GlyT2 and hSERT; DPG3 forms long-lived contacts with two replicates of dDAT and hSERT and all replicates of GlyT2. The lipids with only a glucosylceramide headgroup form fewer sustained contacts with each transporter than is suggested by their high enrichment. Although enriched around all transporters in all replicates, DPGS forms long-lived contacts with only two replicates of GlyT2 compared to all replicates of dDAT and hSERT. Similarly, despite being enriched around all replicates of hSERT and one replicate of GlyT2, PNGS forms no long-lived interactions these transporters in any replicates. Two trajectories of dDAT are enriched in PNGS, and the transporter forms long-lived interactions in one. POGS forms long-lived contacts with hSERT in one replicate, and none with any other transporter. The combined depletion-enrichment and contact analyses suggest that GS lipids with polysaccharide headgroups are enriched around SLC6 transporters when they form long-lived contacts, but those with glucosylceramide headgroups are more mobile and form more transient interactions with each transporter.

SLC6 transporter interactions with GS lipids are localized to particular structural regions, but not to specific amino acid residues (Figure 7). Glycosphingolipids sustain several interactions with all three transporters that persisted for almost an entire 10 μs replicate simulation (99% of the 10 μs simulation in hSERT; 95% in GlyT2; 97% in dDAT). These contacts occur mostly with the glycosphingolipid headgroup and the great majority lie in the loop regions of the protein (Figure 5b). GS lipids most commonly interact with the EL2 region in all three transporters. Here contacts with EL2 account for 41, 48 and 39% of the total GS interactions with hSERT, GlyT2, and dDAT, respectively. Another 14% of the interactions with dDAT and hSERT occur in EL3, but these only account for 6% of interactions with GlyT2. Instead, EL6 is the second most frequently occurring loop interaction for GlyT2 (13%). However, the frequencies of lipid interaction with particular amino acid residues all fall within 6% of their natural abundance (Figure 6b). Glycine and leucine each comprise 9% of GS lipid contacts with dDAT. In GlyT2, contacts with proline account for 11% of interactions and leucine and tryptophan each comprise 9% of interactions with GS lipids. In hSERT, 16% of the residues forming contacts with GS lipids are leucine. The high frequency of interactions with specific amino acids is likely caused by the abundancy of the amino acid and location within the protein, rather than through specific contacts.

**Figure 7.**
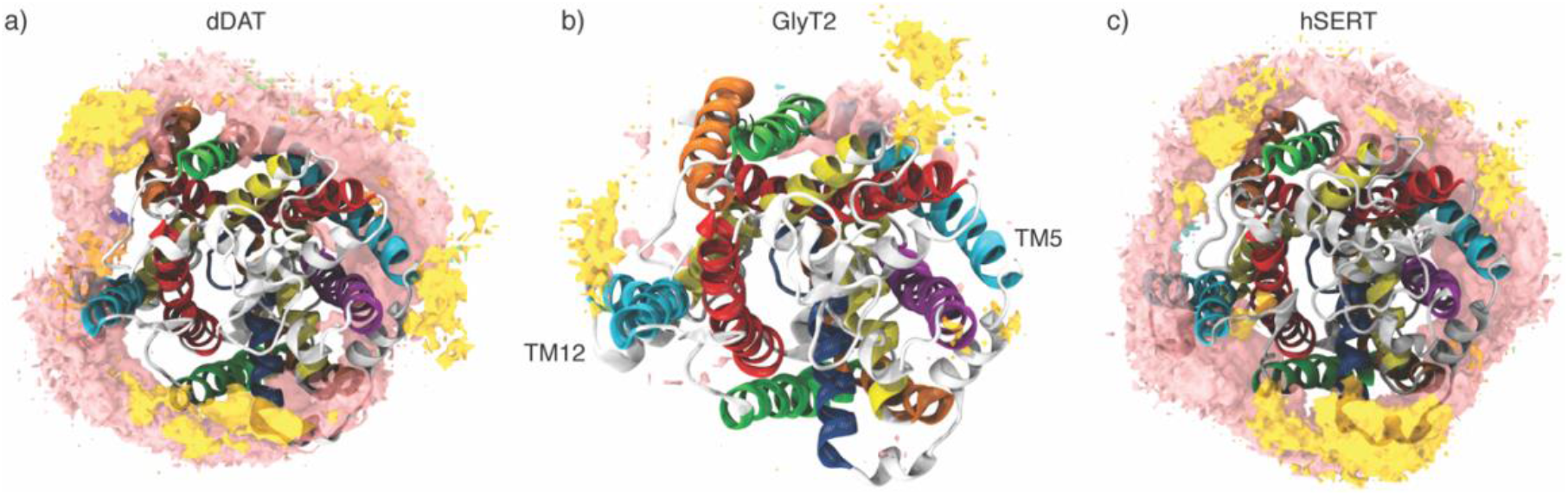
Top-down view of density of lipid groups around a) dDAT, b) GlyT2, and c) hSERT. Surfaces represent spaces with at least 0.005 A^−3^ density. Proteins are aligned. High-density sites are represented in by pink: cholesterol; yellow: GS; and orange: PE. Lower density sites are represented by grey: DAG (PADG); green: PC; blue: PI; and purple: PS. SM lipids are too low-density to be visible.

#### Phosphatidylinositols interact specifically with dDAT, GlyT2, and hSERT, but interactions vary in number and location between the transporters

Phosphoinisotol (PI) lipids comprise 5 mol % of the intracellular leaflet of the neuronal plasma membrane and occur as monounsaturated phosphatidylinositol (POPI), polyunsaturated phosphatidylinositol (PUPI, PAPI and PIPI), and polyunsaturated phosphatidylinositol mono, bi or triphosphate (PAP1, PAP2 and PAP3). As a group, PI lipids are enriched around all three SLC6 transporters investigated (DEI = 3.130 ± 0.342). However, the quantity, length, and location of these interactions varies greatly between each SLC6 transporter, and PI lipids form significantly more sustained contacts with hSERT than the other SLC6 transporters (Figure S6). PI lipids form twice as many persistent interactions (>40% of the simulation time for any 10 μs replicate) with hSERT (76 contacts) than dDAT (34 interactions) or GlyT2 (36 interactions). More of these interactions are long-lived (persisting for ≥ 70% of single 10 μs replicate simulation) in hSERT (11 interactions) and GlyT2 (8 interactions) than in dDAT (1 interaction). Despite the lower abundance of PI lipids in the membrane compared to PE lipids (5 mol% vs 22 mol% of the intracellular leaflet), hSERT forms more long-timescale interactions with PI lipids more than PE lipids. This is not the case for dDAT and GlyT2, where interactions with PI lipids occur less frequently than those with PE or GS lipids.

Only one molecule of the phosphorylated phosphoinisitol lipid, PAP2, exists in the membrane, but it forms persistent interactions with hSERT in all three 10 μs replicates. PAP2 only interacts with GlyT2 in one 10 μs replicate; however, GlyT2 interacts with another phosphorylated phosphoinisitol, PAP1, in a second replicate. Notably, only one phosphorylated lipid (PAP1 or PAP2) interacts with GlyT2 or hSERT at any point in time. Phosphorylated lipids do not form any persistent contacts with dDAT in any of the simulations. PUPI is the most abundant PI lipid (comprising 1 mol% of the membrane), and forms persistent contacts with dDAT and hSERT in all three 10 μs replicates, and with GlyT2 in two of the three replicates. As the lipid population decreases for the different non-phosphorylated PI lipids, more sporadic interactions are observed. The next most abundant species, POPI and PAPI, each comprise 0.6% the membrane. Both POPI and PAPI form persistent contacts with dDAT in two of the three 10 μs replicates, and with hSERT in one 10 μs replicate. No interaction was observed between POPI and GlyT2, while PAPI forms persistent contacts with GlyT2 in one 10 μs replicate simulation. PIPI interacts with hSERT for >40% of the 10 μs simulation time in one replicate simulation but does not interact with GlyT2 or dDAT. The enriched DEI profiles combined with low contact frequencies of each lipid species suggest that there is more exchange with the non-phosphorylated PI lipids than the phosphorylated PI lipids. Indeed, around half of the contacts that occur for >40% of the simulation time in each replicate involve more than one individual PI lipid.

PI lipids interact most commonly with helical regions in dDAT (71% helical regions), while the interactions with GlyT2 and hSERT are more equally distributed across helical and loop regions (58% helical and 42% loop regions). Contacts with the TM5 helix are the most common, comprising 26% of the interactions with PI lipids for dDAT, 17% for GlyT2, and 20% for hSERT (Figure 5c and Figure 8). Each transporter also has unique interaction sites. PI lipids form contacts with dDAT in the region of TM4 (12%), TM7 (12%) and TM11 (18%). For GlyT2, these PI lipid contacts primarily occur at the N-terminus (39%), followed by TM9 (17%), TM5 (14%), and TM4 (14%). In hSERT, 20% of PI contacts occur in the IL2 region. Regardless of the transporter, both the lipid tail (53-60%) and PI lipid headgroup (40-47%) are involved in these interactions.

**Figure 8.**
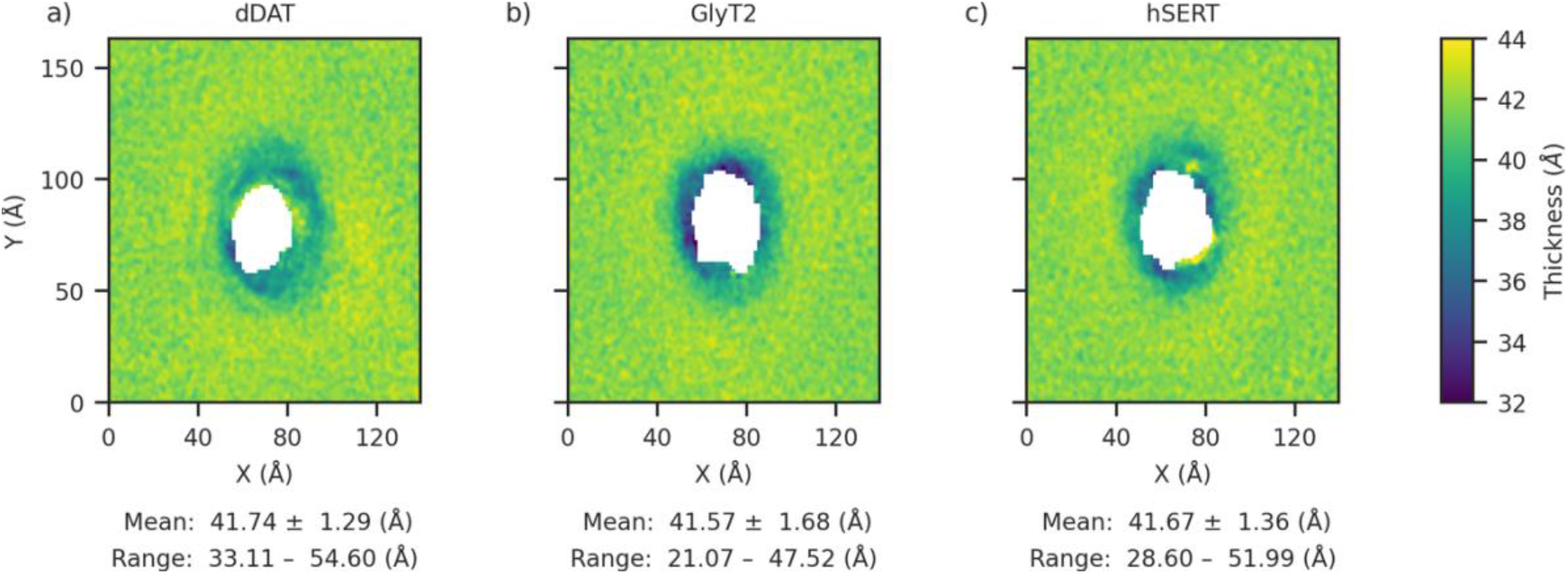
Membrane thickness across the x-y plane of the neuronal membrane embedded with a) dDAT; b) GlyT2; and c) hSERT. Proteins are not aligned to each other. The colour bar is truncated and does not represent the full range of values. Average and standard deviation of the thickness is shown below each plot, followed by the minimum and maximum values.

Figure 6c shows that the specific amino acids in contact with PI lipids also varies between transporters. In hSERT, 24% of all amino acid/PI interactions are with lysine, while in GlyT2 lysine forms 22% of the interactions with PI lipids. This is less so in dDAT, where lysine forms only 15% of the PI interactions, and interactions are also spread across phenylalanine (18%), tryptophan (15%), and arginine (15%). As lysine only comprises between 2.4 and 4% of residues in each protein, these contacts occur far more frequently than may be expected from random interactions. The enrichment of interactions with cationic amino acids is consistent with the PI lipids forming electrostatic interactions with the lipid headgroup. This has been previously reported to be important for stabilizing PI interactions with proteins [40]. In GlyT2, 17% of PI lipid interactions are formed with alanine. In contrast, PI lipid interactions with alanine are not observed in dDAT or hSERT. As 7.5–8.8 % of each transporter is alanine, the preference of PI lipids for interacting with alanine in GlyT2 and avoidance of alanine in dDAT and hSERT is another example of the different interaction profiles of each transporter. The interactions with alanine in GlyT2 occur in TM4 and with Ala190 in the N-terminus; in hSERT, the N-terminus and TM4 interactions involve tryptophan, glycine, lysine, valine, and isoleucine. In dDAT, the N-terminus and TM4 interactions are with arginine, tryptophan, phenylalanine, and serine.

The binding of phosphatidylinositol (4,5)-bisphosphate to the N-terminus (Lys3 and Lys5) of hDAT has been shown to support dopamine release in response to amphetamine [40]. PI lipids form contacts with the N-terminus of all three transporters; however, these interactions do not exclusively occur with phosphatidylinositol bisphosphate lipids. Rather, most interactions with the N-terminus occur with non-phosphorylated PI lipids, although we also observe contacts with phosphatidylinositol monophosphate and phosphatidylinositol bisphosphate lipids. Additionally, while previous contacts were reported with the Lys3 and Lys5 of hDAT, the crystal structure has 24 amino acids truncated from the N-terminus, thus different amino acids form the basis of the interactions. In hSERT, the N-terminus residues (Trp82, Gly83 and Lys84) largely interact with POPI. In dDAT (Arg27 and Trp30), the interactions are with PUPI. Several N-terminus residues of GlyT2 (Lys189, Ala190, Arg191, Gly192, Asn193 and Trp194) interact with PAP2, PUPI, and PAP1.

#### Phosphatidic acids, sphingomyelins, diacylglycerols and ceramides form few sustained contacts with SLC6 transporters

The interactions of several lipid classes with hSERT, dDAT and GlyT2 are distinctly depleted (Figure 4) or transient. Sphingomyelins (SM) comprise 9% of the extracellular leaflet and 2% of the intracellular leaflet but form few contacts with each transporter. SM lipids are depleted around all proteins in both leaflets (DEI_extracellular_ = 0.490 ± 0.114; DEI_intracellular_ = 0.374 ± 0.084) and do not form direct contacts with any of the SLC6 transporters for >50% of each 10 μs replicate simulation. The low concentration lipids, CER (ceramide, 1 DPCE in each leaflet) and PA (1 PAPA in the membrane), form few contacts with each transporter; DAG (1 PADG in each leaflet) has transient contacts with each transporter. None of these lipids formed systematic contacts with any transporter over multiple replicate simulations. However, due to the low number of these lipids within each leaflet, any localization around the transporter was magnified in the DEI analysis. Despite there being only two molecules of DPCE in the neuronal membrane system, one molecule of DPCE forms a direct contact with dDAT in one 10 μs replicate. This contact lasted for 75% of the simulation time, in which DPCE was bound in the region between TM2, TM6 and TM11 (DEI_extracellular_ = 3.372; DEI_intracellular_ = 7.904). In the remaining two replicate simulations of dDAT, it is otherwise very depleted around the transporter (Table S2). DPCE was also depleted around and hSERT in all replicate simulations (Table S2). The one molecule of PAPA in the nueronal membrane does not form contacts with any of the three transporters for over 40% of the time in any 10 μs replicate simulation and is depleted around all three SLC6 transporters (DEI_intracellular_ = 0.493 ± 0.370). In contrast, despite the presence of only two molecules of PADG in the neuronal membrane, PADG is significantly enriched around each of the three transporters in both leaflets (DEI_extracellular_ = 5.324 ± 2.260; DEI_intracellular_ = 3.620 ± 1.643). Nonetheless, it forms persistent contacts only in single 10 μs replicates of GlyT2 and hSERT, and no persistent contacts with dDAT. PADG flips between leaflets, and the glycerol groups of PADG interact directly with GlyT2 and hSERT near the interleaflet interface. Specifically, a single PADG lipid interacts for 67% of a single 10 μs GlyT2 replicate simulation in the region between TM2 and TM7. PADG also forms contacts with hSERT for 68% of the simulation time in one replicate, predominantly in the region of TM10 but also with TM4, TM6 and TM7. This indicates that PADG selectively aggregates around each transporter but does not form sustained contacts with any single region. Overall, the rare lipids sphingomyelins, phosphatidic acids, diacylglycerols and ceramides did not form persistent contacts with any of the three neurotransmitter transporters examined.

### Cholesterol binds to SLC6 transporters at specific sites on each protein

Cholesterol makes up 47% of the neuronal plasma membrane and is known to play a key role in neurotransmission, and the activity of DAT [41,42] and SERT [43,44]. GlyT2 is modulated by different concentrations of cholesterol in the membrane [45]. Bound cholesterol has been found at the interface of TM1a, TM5, and TM7 (which form the CHOL1 binding site) in all structures of dDAT [17,46,47], while the cholesterol analogue, cholesteryl hemisuccinate, is bound at the interface of TM2 and TM7 (at the CHOL2 binding site) in a subset of these structures [17,47]. A potential third cholesterol binding site (CHOL3) has been identified on TM12 of the hSERT crystal structure [18]. In addition, previous coarse-grained simulations of hSERT and dDAT in POPC/CHOL membranes located another three binding sites around TM4, TM5, and TM8 (CHOL4); TM9 and EL2 (CHOL5); and TM10, TM11, and TM12 (CHOL6) [48]. There are numerous occurrences of cholesterol-binding motifs (CRAC and CARC) in each protein. Specifically, 4 CRAC motifs and 10 CARC motifs are present in dDAT; 5 of each CRAC and CARC motifs reside on GlyT2; and 6 CRAC and 5 CARC motifs are present on hSERT. While these motifs can be predictive of cholesterol binding [49], previous simulations have found that most of these motifs are not near major binding sites [48]. Similarly, we observe that cholesterol aggregates preferentially around two particular cholesterol-binding motifs, instead of binding equally to every motif present.

Across our simulations, cholesterol is slightly depleted around all three SLC6 transporters in the neuronal membranes (DEI_extracellular_ = 0.919 ± 0.028, DEI_intracellular_ = 0.919 ± 0.039, Figure 4). However, the remaining interactions with the protein are highly localised to certain regions (Figure S7). Cholesterol forms 462 persistent contacts (lasting >40% of each 10 μs replicate simulation) with dDAT, 483 with GlyT2, and 465 with hSERT. Of these, 109 contacts in dDAT are long-lasting and are sustained >90% simulation time for each 10 μs replicate, compared to 122 contacts in GlyT2, and 104 in hSERT. As expected, long-lasting contacts are formed almost exclusively in helical regions (96% long-lasting contacts in dDAT; 99.2% in GlyT2; 99.0% in hSERT). A substantial number of the long-lasting contacts are formed with residues within 6 Å of previously identified binding sites, particularly those present in the crystal structures. In the case of dDAT, 25% of the long-lasting cholesterol contacts are formed with residues within 6 Å of crystallographic cholesterol at binding sites CHOL1 and CHOL2, another 4% of contacts are formed with residues around CHOL3. In GlyT2, 20% long-lasting contacts are also formed at regions corresponding to the crystallographic binding sites: while cholesterol does not occupy the CHOL1 binding site of GlyT2 significantly (5% long-lasting contacts), it forms many long-lasting contacts with residues at the CHOL2 and CHOL3 sites (15% contacts). In the case of hSERT, 18% of the long-lasting cholesterol contacts are at the crystallographic binding sites; while cholesterol is found at CHOL1 and CHOL2 and with TM12 in general but cholesterol forms only one long-lasting contact at CHOL3.

Cholesterol is also found to occupy locations corresponding to the binding sites previous identified in simulation by Zeppelin *et al*. [48]. In dDAT, cholesterol forms contacts at binding sites CHOL4 and CHOL5; however, it does not have high occupancy of the CHOL6 site. In fact, contacts with TM10 comprise only 2% the long-lasting contacts formed with dDAT. The greatest proportion of long-lived contacts are formed with TM5, both at the nearby binding sites CHOL1 and CHOL4 and with the helix itself. Although a CARC motif is present on TM5, most sustained contacts are not formed with residues of the CARC motif, but at the extracellular end of the helix instead. In GlyT2, cholesterol is present at CHOL4 and CHOL5 but forms substantially more contacts around CHOL6, with 28% of long-lasting contacts formed with residues around CHOL6. In hSERT, cholesterol again forms substantial contacts at CHOL6 (33% long-lasting contacts) and CHOL4 (16% long-lasting contacts), but cholesterol does not associate significantly with the transporter at CHOL5.

While site-specific binding of cholesterol is observed in all three SLC6 transporters, CRAC and CARC motifs comprise a high percentage of the protein sequence (~25%), and cholesterol does not preferentially bind to all these cholesterol-binding motifs in dDAT, hSERT and GlyT2. Instead, the majority of cholesterol binding at cholesterol-binding motifs is targeted to two specific cholesterol-binding motifs (CRAC motif on TM4 and IL2 and CARC motif on TM11). Specifically, cholesterol binds to CRAC motif on TM4 and IL2 on all three transporters; and to residues of the CARC motif on TM11 on dDAT and GlyT2. This latter motif is not present on hSERT. In total, another 11% long-lasting contacts are formed with the specified CRAC and CARC motifs on dDAT, compared to 15% long-lasting contacts with GlyT2, and 9% long-lasting contacts with hSERT. The lack of the CARC motif on TM11 on hSERT, but substantial binding to a binding site (CHOL6) that includes TM11, is further indication that cholesterol-binding motifs are not a prerequisite for cholesterol-binding. This is supported by the binding of cholesterol to specific motifs rather than all CARC and CRAC motifs.

Over all three SLC6 transporters, cholesterol binds to Ile, Leu, and Phe up to 10% more than the natural abundance of these amino acid in dDAT and hSERT (Figure 6d). In GlyT2, contacts between CHOL and Ile or Leu occur up to 6% more than their natural abundance. This is consistent with branched amino acid (e.g., Ile and Leu) associating with the β-face of cholesterol through van der Waals interactions [50], and aromatic side-chains forming CH-π interactions with the α-face of cholesterol [51]. In all three transporters, contacts with Gly occur up to 7% less than its natural abundance, while all other amino acids of the transporters interact with CHOL within 4% of their occurrence. In dDAT, TM5 has the greatest proportion of extremely long-lived contacts (21%) (Figure 5d). In hSERT, the most long-lived contacts are formed with residues in TM11 (22%) and TM5 (21%). In GlyT2, long-lived cholesterol contacts with TM5 and TM11 only comprise 11% each of the total long-lived contacts. Instead, most cholesterol interactions for GlyT2 occur with TM12 (18%) and TM8 (12%). Overall, while similar binding sites are observed for each SLC6 transporter, cholesterol binds preferentially at different sites for each protein: CHOL1 is the most populated site in dDAT, compared to CHOL6 in hSERT, and both CHOL3 and CHOL6 in GlyT2.

### The presence and identity of embedded SLC6 transporters alter some membrane properties

To evaluate the effects of each SLC6 transporter on the neuronal membrane biophysical properties, we calculated the membrane thickness, area per lipid, lipid lateral self-diffusivity and distribution of lipid species for each neurotransmitter transporter system. Due to the asymmetric composition of the neuronal membrane, results are determined for each leaflet individually.

The average membrane thickness does not vary significantly when the three SLC6 transporters are embedded in the membrane. The dDAT and hSERT neuronal membranes both have a thickness of 41.7 ± 1.3 Å, while the thickness of the GlyT2 neuronal membrane is 41.6 ± 1.7 Å (Table 1). These values are consistent with the membrane thickness derived from previously published simulations of neuronal membrane (41.49 ± 0.22 Å) [13] in the absence of membrane proteins, suggesting that the presence of the transporter does not significantly affect this property. The thickness is not a uniform property across the membrane (Figure 8) but varies by proximity to the embedded transporter. In all cases, the membrane is thinner closer to the protein-lipid interface. In the neuronal systems, the maximum membrane thickness is 47.5, 52.0 or 54.6 Å for GlyT2, hSERT and dDAT, respectively (Table 1). Similarly, the same trend remains when the thinnest point in the membrane is considered with GlyT2 having lowest minimum membrane thickness (21.1 Å), followed by hSERT (28.6 Å) and dDAT (33.1 Å). This is despite all three neuronal membranes having the same chemical composition. It is unclear why the membrane thickness for dDAT embedded in the neuronal membrane is higher than that of GlyT2 and hSERT. Polyunsaturated lipids are known to lower membrane thickness, but the lipid ring around dDAT is the most enriched in polyunsaturated lipids of all the SLC6 transporters and is slightly depleted in monounsaturated lipids in the extracellular leaflet.

**Table 1.** Neuronal membrane thickness and area per lipid of each leaflet with and without SLC6 transporters

| SLC6 transporter | Average thickness (Å) | Minimum thickness (Å) | Maximum thickness (Å) | Area per lipid (APL) |  |
| --- | --- | --- | --- | --- | --- |
|  |  |  |  | Intracellular leaflet (Å<sup>2</sup>) | Extracellular leaflet (Å<sup>2</sup>) |
| dDAT | 41.7 ± 1.3 | 33.1 | 54.6 | 51.8 ± 0.3 | 49.6 ± 0.3 |
| GlyT2 | 41.6 ± 1.7 | 21.1 | 47.5 | 51.8 ± 0.3 | 49.6 ± 0.3 |
| hSERT | 41.7 ± 1.4 | 28.6 | 52.0 | 51.9 ± 0.3 | 49.6 ± 0.3 |
| No protein<sup>a</sup> | 42.2 ± 0.2 | 41.6 | 42.9 | 56.4 ± 0.8 | 50.1 ± 0.7 |
| No protein<sup>b</sup> | 40.6 ± 0.02 | -<sup>c</sup> | -<sup>c</sup> | 48.5 ± 0.1 | 46.0 ± 0.1 |

<sup>a</sup>Data taken from Wilson *et al.*, 2020 [13] and recalculated with *g\_thickness* to be consistent with the present work. <sup>b</sup>Data taken from Ingólfsson *et al.*, 2017 [11]. <sup>c</sup>Not provided.

**Table 2.** POPC/CHOL model average membrane thickness and area per lipid with and without SLC6 transporters

| SLC6 transporter | Average thickness (Å) | Minimum thickness (Å) | Maximum thickness (Å) | Area per lipid (APL) |  |
| --- | --- | --- | --- | --- | --- |
|  |  |  |  | Intracellular leaflet (Å<sup>2</sup>) | Extracellular leaflet (Å<sup>2</sup>) |
| dDAT | 41.9 ± 0.2 | 27.8 | 42.9 | 51.8 ± 0.3 | 49.6 ± 0.3 |
| GlyT2 | 41.2 ± 0.9 | 30.1 | 45.3 | 51.8 ± 0.3 | 49.6 ± 0.3 |
| hSERT | 41.2 ± 0.9 | 27.5 | 46.1 | 51.9 ± 0.3 | 49.6 ± 0.3 |
| No protein<sup>a</sup> | 40.8 ± 0.1 | 40.5 | 41.0 | 56.4 ± 0.8 | 50.1 ± 0.7 |

<sup>a</sup>Data taken from Wilson *et al.*, 2020 [13] and recalculated with *g\_thickness* to be consistent with the present work.

The area per lipid (APL) is related to the packing, phase, and fluidity of the membrane [52,53]. The average APL is not sensitive to which protein is embedded, but differs between leaflets. Table 1 shows the average APL of the extracellular leaflet is lower than that of the intracellular leaflet. However, these values lie within the range previously reported for simulations of neuronal membranes in the absence of protein (Table 1) [11,13], suggesting that the presence of the embedded SLC6 transporter does not significantly affect the APL. One factor that affects the APL is the proportion of polyunsaturated lipids in the membrane, as polyunsaturated lipids have larger areas per lipid (Figure S8) [54]. In the neuronal membrane, 36% the intracellular leaflet is comprised of polyunsaturated lipids, compared to only 23% the extracellular. In addition, the relative enrichment of low APL lipids in the extracellular leaflet suggests a possible basis for the difference in areas per lipid between the leaflets. Of the lipid species present, glycosphingolipids (36.8–37.5 Å^2^) and sphingomyelins (45.7–48.3 Å^2^) have the smallest areas per lipid (Figure 9) and are predominantly found in the extracellular leaflet (Table S1), explaining the lower APL of the extracellular leaflet.

**Figure 9.**
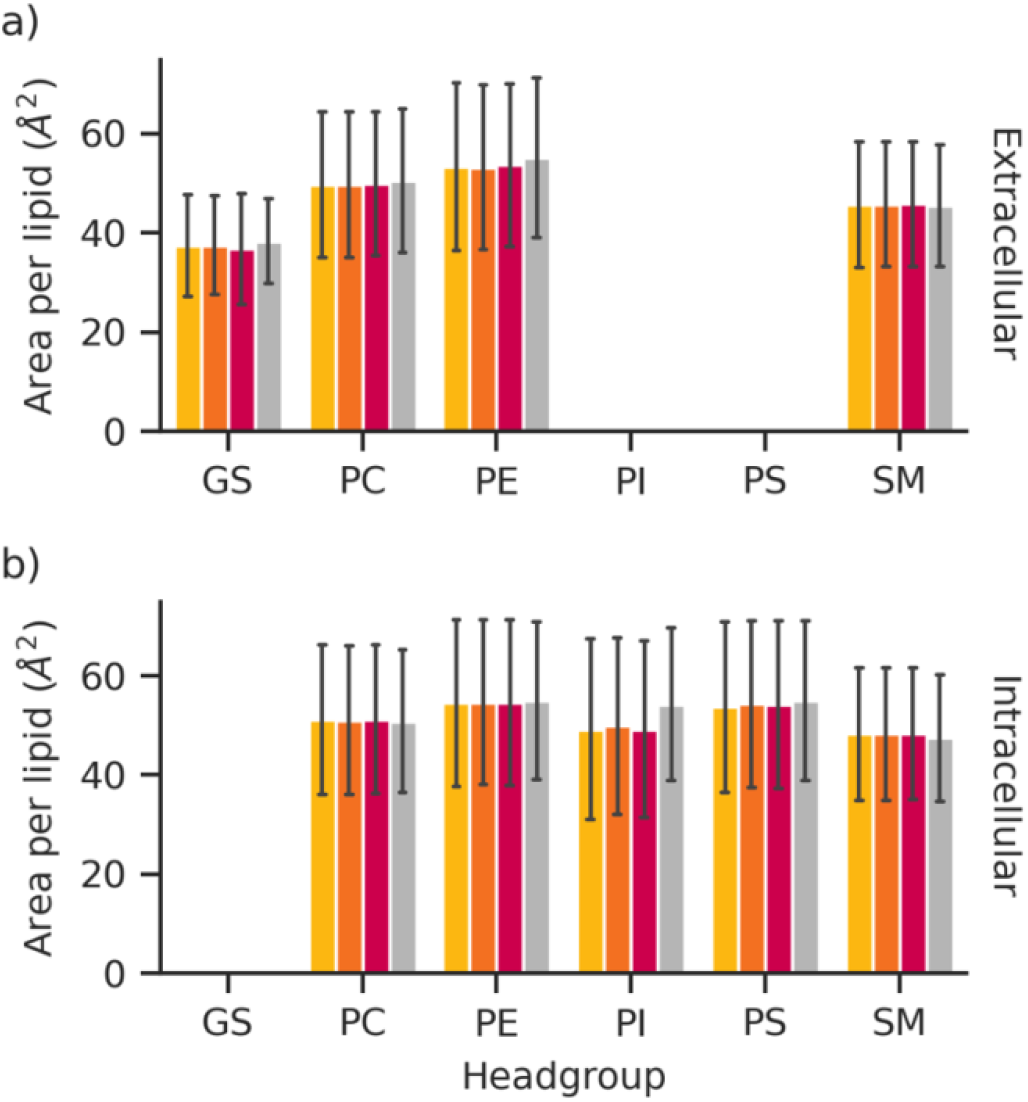
Area per lipid for each lipid class in the neuronal membrane, for a) the extracellular leaflet; and b) the intracellular leaflet. The standard deviation is represented by the error bars. Yellow: dDAT. Orange: GlyT2. Red: hSERT. Grey: no protein [13].

Regardless of which SLC6 transporter is present, the lateral self-diffusivity of the lipids in the extracellular leaflet is slower than in the intracellular leaflet. This is consistent with previous observations for the neuronal plasma membrane without protein [13]. The lateral self-diffusivity of GS and PI lipids, which form persistent specific contacts with the transporter, are not affected by the presence of the transporter (0.3-0.4 × 10^−7^ and 1.1-1.4 × 10^−7^ cm^2^/s, respectively; Figure 10). Furthermore, the lateral self-diffusivity of PE and PS lipids, which form non-specific contacts with the transporter, are within 0.1 × 10^−7^ cm^2^/s of the neuronal membrane for both the extracellular and intracellular leaflets (0.8-0.9 and 1.4-1.5 × 10^−7^ cm^2^/s, respectively) [13]. When embedded dDAT is present, the lateral self-diffusivity of PC lipids is within 0.1 × 10^−7^ cm^2^/s of the neuronal membrane (0.8 and 1.4 × 10^−7^ cm^2^/s for the extracellular and intracellular leaflets, respectively). However, in the presence of hSERT, the lateral self-diffusivity of PC lipids is increased by 0.2 × 10^−7^ cm^2^/s for both the extracellular and intracellular leaflets compared to the neuronal membrane. When GlyT2 is present, the lateral self-diffusivity of PC lipids is increased by 0.1 or 0.3 × 10^−7^ cm^2^/s for the extracellular and intracellular leaflets, respectively compared to the neuronal membrane. Finally, there is a significant increase in the lateral self-diffusivity of SM lipids in the extracellular leaflet of the neuronal membrane in the presence of all three SLC6 transporters (0.8 × 10^−7^ cm^2^/s), when compared to the membrane alone (0.5 × 10^−7^ cm^2^/s). Conversely, in the intracellular leaflet, the rate of SM lateral self-diffusivity is lower when transporter is present (1.0-1.2 × 10^−7^ cm^2^/s) than in the isolated membrane (1.3 × 10^−7^ cm^2^/s). Overall, the changes in lateral self-diffusivity does not appear to be correlated to the propensity of the lipid to form interactions with the transporters.

**Figure 10.**
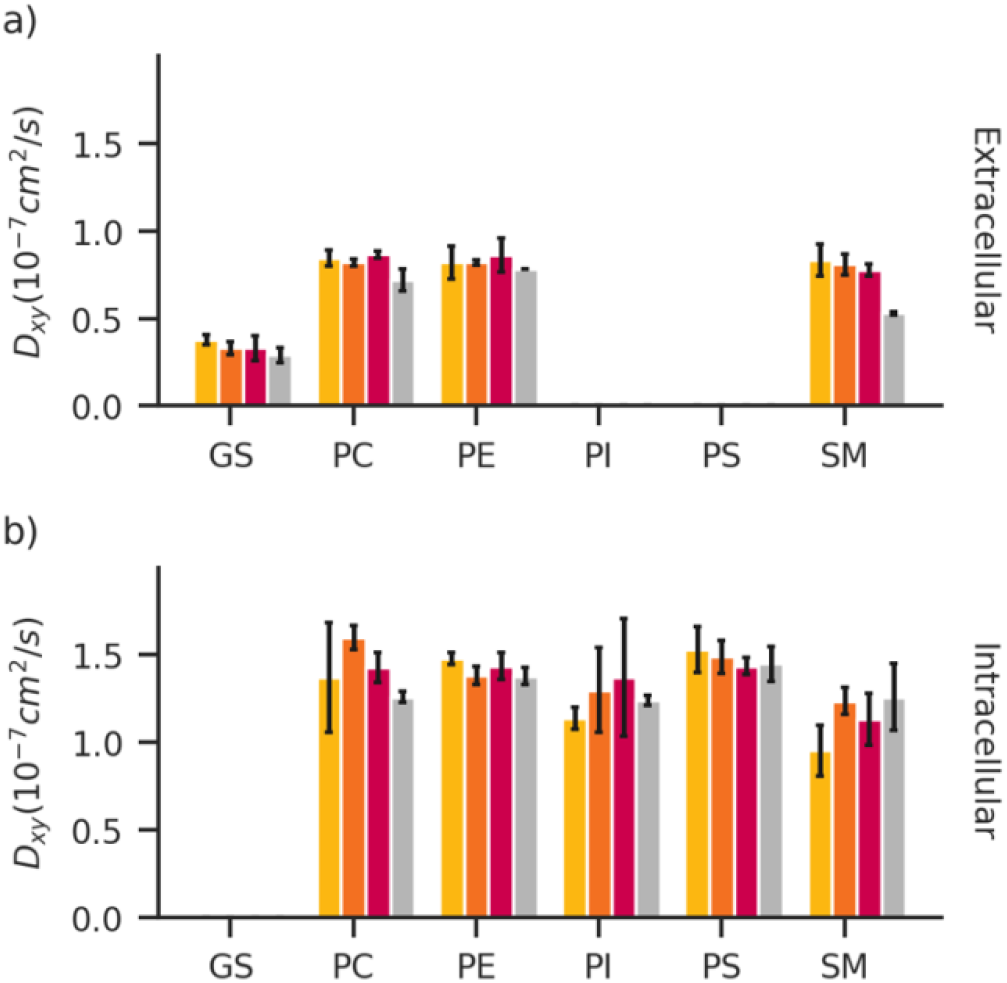
Self-diffusivity of lipid classes in the neuronal membrane, for a) the extracellular leaflet; and b) the intracellular leaflet. The standard error from the mean is represented by the error bars. Yellow: dDAT. Orange: GlyT2. Red: hSERT. Grey: no protein [13].

The same pattern of domain formation occurs in the neuronal membranes regardless of which SLC6 transporter is present (Figure 11). Specifically, CHOL is equally distributed throughout the membrane and PC lipids are slightly enriched around SM lipids (1.3) but otherwise equally distributed. PE lipids are enriched around PE (1.2-1.3), PS (1.6) and PI (1.7) but depleted around GS lipids (0.5). PS lipids are highly enriched in the vicinity of PS and PI lipids (2.0 and 2.1, respectively) and are depleted around SM (0.5). SM and GS are enriched around each other with significant rafting of GS observed (3.2-3.4), while SM-SM and GS-SM clustering is enriched to 1.6. PI lipids also raft (2.2-2.3) and are depleted around SM (0.4). In general, the domain formation is similar to that observed in the neuronal membrane in the absence of the embedded transporters [13] with exception to PI domain formation that occurs less in the presence of the transporter and GS domain formation that occurs more in the presence of the transporters. Since both PI and GS lipids form persistent interactions with the SLC6 transporters this indicates that these interactions are changing how the lipids are positioned in the membrane. Indeed, transporters have been shown to interact with GS rafts within membranes [55].

**Figure 11.**
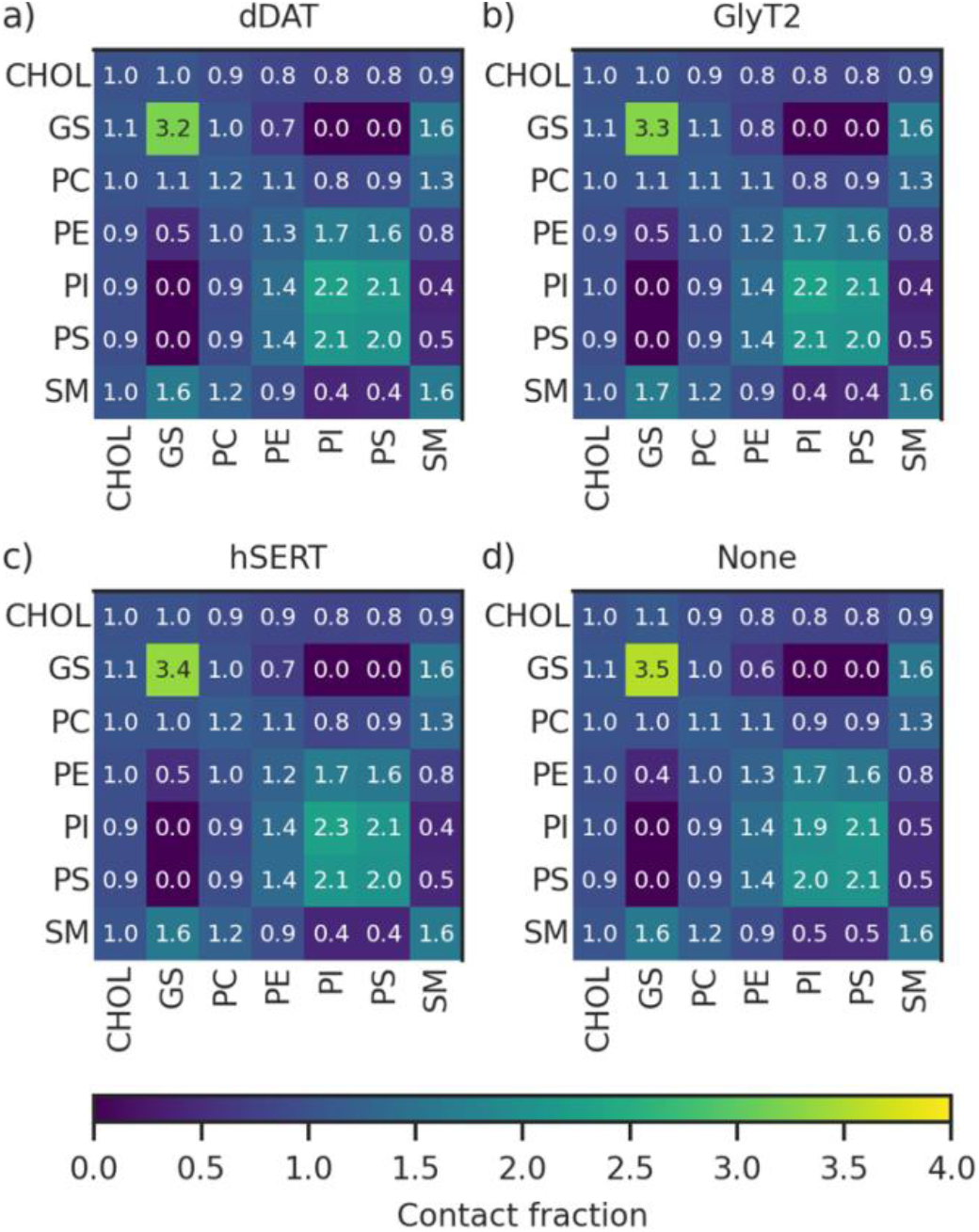
The average lipid contact fractions of the neuronal membrane, by head group, for membranes with embedded: **a)** dDAT, **b)** GlyT2, **c)** hSERT, and **d)** no protein [13].

Cholesterol flip-flop between leaflets occurs at a faster rate (3.5 ± 0.1 × 10^6^ s^−1^) in the neuronal membrane with the transporter embedded than the rates reported for corresponding membranes without embedded proteins (2.3 × 10^6^ s^−1^) but slower than the rate of cholesterol flip-flop (4.820 × 10^6^ s^−1^) in the much larger neuronal membrane reported by Ingolfsson *et al.* [11]. Absolute flip-flop rates are notoriously hard to compare as they change depending on the time interval of the frames included for analysis (flip-flop events can be missed within large time intervals). The reduction in flip-flop in the neuronal membrane reflects previous research showing that cholesterol flip-flop decreases as cholesterol content is increased, and increases with bilayer polyunsaturation. [56] The flip-flop rate does not change significantly when different transporters are embedded in the membrane.

In general, the properties of the neuronal membrane appear to vary within the same range as those of the neuronal membrane without a transporter embedded, indicating that the presence of a protein does not significantly alter properties. The exceptions are the lipid lateral self-diffusivity, which is increased for SM and PC lipids, and PI and GS domain formation, which are altered by interactions with the SLC6 transporters. Comparing membrane properties when different transporters are embedded found that the thickness profile of the membrane over distance from the transporter, varied between different SLC6 transporters. Lipid lateral self-diffusivity also varied between proteins for PC and SM lipids. The area-per-lipid, cholesterol flip-flop rate, and domain formation, however, remained constant between different transporters.

### Effect of Complex membrane composition

We compared membrane properties of the neuronal membrane and the model membrane to investigate the effect of the SLC6 transporters on properties that are affected by membrane composition. For the neuronal membrane systems, dDAT has the greatest membrane thickness (41.9 ± 0.2 Å), while the GlyT2 and hSERT systems have a membrane thickness of 41.2 ± 0.2 Å. There is less variation in the thickness of the POPC/CHOL model membrane systems compared to the neuronal membrane, with membrane thickness ranging from 39.8–40.1 Å. The presence of protein does not significantly influence average membrane thickness. The average membrane thickness for both the neuronal and POPC/CHOL membranes are consistent with values derived from previously published simulations of neuronal (41.5 ± 0.2 Å) and POPC/CHOL (41.0 ± 0.2 Å) membranes without SLC6 transporters embedded [13]. As noted previously, the thickness of the membrane varies with proximity to the protein in the neuronal membrane. While the maximum thickness of the membrane varied for each transporter in the neuronal membrane, the maximum membrane thickness is 40.8 Å regardless of the embedded transporter in the model POPC/CHOL systems. The minimum membrane thickness occurs closest to the protein and is 34.9–35.5 Å. Overall, the neuronal membrane systems are 1–2 Å thicker than the model membrane systems (Figure S9).

POPC has a greater area per lipid in the POPC/CHOL simulations (58.8 ± 14.7 Å^2^) than the neuronal membrane systems (48.7 ± 13.4 Å^2^), regardless of which SLC6 transporter is embedded. The higher concentration of cholesterol in the neuronal membrane is likely to play a role in the difference in properties between the neuronal and POPC/CHOL membranes. Cholesterol has been shown to induce the ordering of lipid fatty acid tails, which would increase membrane thickness and correspondingly reduce the area per lipid [57]. Experimentally determined APLs of simplified membranes show decreasing average areas as the concentration of cholesterol increases [13]. The average APL for a 50% CHOL/50% POPC membrane has been determined to be 45.1 ± 0.9 Å^2^ [58], and the average APL for a 40% CHOL/60% POPC membrane to be 46 Å^2^ [59]. In addition, we find that polyunsaturated lipids have a significantly larger APL (53.6 ± 17.2 Å^2^) compared to monounsaturated (45.6 ± 13.5 Å^2^) and saturated (47.0 ± 12.5 Å^2^) lipids in the neuronal membrane, supporting the supposition that the increased ordering is one reason for the difference in APL.

Cholesterol flip-flop between leaflets occurs at a slower rate (3.5 × 10^6^ s^−1^) in the neuronal membrane than the POPC/CHOL membranes (4.5 × 10^6^ s^−1^). As with the neuronal membrane, flip-flop in the simple membrane does not vary significantly between different proteins and occurs substantially faster than a corresponding model membrane without the transporter embedded (POPC/CHOL, 3.7 × 10^6^ s^−1^) [13]. The reduction in flip-flop in the neuronal membrane reflects previous research showing that cholesterol flip-flop decreases as cholesterol content is increased, and increases with bilayer polyunsaturation [56].

Unlike the neuronal membrane, the properties of the simple POPC/CHOL model do not vary significantly between the different SLC6 transporters. It is likely that a diversity of lipids is necessary for individual lipid-protein interactions to have a meaningful effect on average membrane properties. Furthermore, as previously discussed many persistent interactions are formed between PE, PI and GS lipids in the neuronal membrane that will not be accounted for in a simplified POPC/CHOL membrane system.

### The local lipid environment of SLC6 transporters in the neuronal membrane differs from the plasma membrane

The composition of the neuronal membrane differs from the average plasma membrane in several ways. The neuronal membrane contains significantly more cholesterol by proportion than the average membrane; cholesterol constitutes 46.8% the neuronal membrane, compared to 30.1% the average model. By tail saturation, the neuronal membrane contains two saturated lipid species (DPPC and DPPS) that together form 4.4% the neuronal membrane; the average plasma membrane, on the other hand, contains only a single saturated species (PPC) that comprises 0.3% the average membrane. Monounsaturated lipids are less abundant in the neuronal membrane, especially the intracellular leaflet. They comprise 13.5% the intracellular leaflet of the neuronal membrane, compared to 23.1% the intracellular leaflet of the average. The proportion of polyunsaturated lipids is also substantially lower in the neuronal membrane (29.5%) than the average plasma membrane (44.5%). By headgroup, the biggest differences in membrane composition are with the PC, PE, and SM lipids. PC and SM lipids are less abundant in the neuronal membrane than the average plasma membrane, particularly in the extracellular leaflet. PC lipids comprise only 24.1% the extracellular leaflet and SM lipids only 9.0% the extracellular leaflet of the neuronal membrane compared to 35.6% and 19.0% the extracellular leaflet of the average membrane, respectively. PE lipids are somewhat more abundant in the extracellular leaflet of the neuronal membrane (11.0% compared to 5.6% of the average membrane) but less abundant in the intracellular leaflet (21.9% compared to 25.7% of the average membrane). As membrane behaviour is highly dependent on membrane composition, it is worthwhile to investigate to what extent a protein “lipid fingerprint” varies between the average and neuronal membranes. As dDAT has been modelled in both the neuronal and plasma membrane, this provides an ideal test case to investigate the effects of composition on the “lipid fingerprint”.

We observe that although the general composition of the lipid annulus around dDAT is similar between the membranes, the proportions differ significantly. For example, polyunsaturated lipids are enriched around dDAT in both membranes. However, a leaflet-averaged DEI value of 4.161 ± 1.342 was calculated for polyunsaturated lipids within a 7 Å radius around dDAT in the average membrane. This indicates markedly higher enrichment of polyunsaturated lipids around dDAT in the average membrane compared to the neuronal membrane (DEI_extracellular_ = 1.391 ± 0.179, DEI_intracellular_ = 1.243 ± 0.034). Polyunsaturated lipids comprise a significantly lower portion of the neuronal membrane (29.5%) than the average plasma membrane (44.5%), which means that similar numbers of polyunsaturated lipids aggregating around the transporter would result in much higher DEI values for the neuronal membrane than the average. This suggests that the difference in membrane composition has resulted in substantially lower numbers of polyunsaturated lipids around dDAT. The behaviour of saturated lipids is also significantly affected, although this may be due in part to their low numbers. Fully saturated lipids are slightly enriched around dDAT (DEI = 1.258 ± 0.218) in the average membrane; they are significantly depleted around dDAT in both leaflets of the neuronal membrane (DEI = 0.466 ± 0.093). Monounsaturated lipids are depleted in the intracellular leaflet (DEI = 0.715 ± 0.149) but neither enriched nor depleted in the extracellular leaflet (DEI = 1.015 ± 0.144). The enrichment or depletion of monounsaturated lipids as a class is not reported by Ingólfsson *et al*. for the neuronal membrane [11].

Differences also occur based on the enrichment of lipids when classified by headgroups as a function of membrane composition. Glycosphingolipids are far more enriched around dDAT in the average membrane (DEI = 6.367 ± 0.285) than the neuronal membrane (DEI = 1.882 ± 0.296), although the proportion of glycosphingolipids in the neuronal membrane (11.0%) is double the proportion in the average membrane (5.6%). PC lipids are less depleted in the neuronal membrane (DEI = 0.706 ± 0.112) than the average membrane (DEI = 0.464 ± 0.047), possibly due to lower abundance in the former. SM lipids are similarly less depleted in the neuronal membrane (DEI = 0.432 ± 0.114) than the average plasma membrane (DEI = 0.175 ± 0.058). As with PC lipids, SM lipids comprise significantly less of the neuronal membrane than the average plasma membrane. There are significant differences in the enrichment profiles of DAG, CER, and PA lipids, but these are likely due to their low population in the neuronal membrane (0.2%, 0.2%, and 0.1% the average membrane respectively; there are 2 molecules each of DAG and one PA lipid in the neuronal membrane). While these groups are also a small part of the average plasma membrane, they are both more numerous in absolute number and comprise proportionally more of the membrane: 41 DAG lipids form 0.6% of the average membrane, 40 CER lipids constitute another 0.6%, and 46 PA molecules form 0.7% of the average membrane.

## CONCLUSION

In this study we investigate how the lipid annulus of three SLC6 transporters differs, both within members of the same class of proteins and in membranes of different composition. We found that the overall lipid fingerprints were similar between each SLC6 transporter when lipids were grouped by saturation and headgroup. Polyunsaturated lipids, which comprise the vast majority of the membrane, are enriched around each transporter. PC and PS lipids form few, non-specific and non-sustained contacts with each transporter. PE, PI and GS lipids are enriched around each transporter, but each class interacted with the SLC6 transporter differently; the tail of PE lipids interacted non-specifically with each transporter, while PI headgroups and tails interact non-specifically with each transporter in similar ratios. In contrast, GS headgroups formed persistent contacts with protein loop regions. However, there were unique differences at the chemical species level in each lipid class with respect to enrichment around the transporter, or number and length of contacts formed. In addition, cholesterol clusters around common binding sites, but binds preferentially at different binding sites for each transporter. We find that the composition of the lipid annulus is heavily dependent on the composition of the membrane. In comparing the composition of the local lipid environment of SLC6 transporters embedded in an average plasma membrane to the neuronal membrane, we found that the enrichment of chemically distinct lipid species around each transporter, and the lipid-protein interactions, were likely to be affected by the proportion of specific lipid species in the membrane. This is particularly true for low concentration species, where only one or two lipids are present in the neuronal membrane.

We also characterized the effect of embedded SLC6 transporters on the biophysical properties of simple and complex membranes. For many metrics, such as the area per lipid and the average thickness, the presence of a protein did not alter the membrane properties when compared to the corresponding protein-free membrane for both the simple and the neuronal membrane. However, properties such as the local membrane thickness and lipid lateral self-diffusivity varied between SLC6 transporters in the neuronal membrane, but no such variation was observed in the simple POPC/CHOL model. This highlights the importance of studying membrane-protein interactions in a membrane model that accurately represents both the complexity and composition of the biological environment.

## Supporting information

Supporting Information Figures 1-10, expanded methods

## ACKNOWLEDGMENTS

This work was supported by the National Health and Medical Research Council Project Grant APP1144429. The research was undertaken with the assistance of resources and services from the National Computational Infrastructure (NCI), which is supported by the Australian Government.

## REFERENCES

[1] K.A. Wilson, L. Wang, H. MacDermott-Opeskin, M.L. O’Mara, The Fats of Life: Using Computational Chemistry to Characterise the Eukaryotic Cell Membrane, Aust. J. Chem. 73 (2020) 85–95. https://doi.org/10.1071/CH19353.

[2] J.S. Allhusen, J.C. Conboy, The Ins and Outs of Lipid Flip-Flop, Acc. Chem. Res. 50 (2017) 58–65. https://doi.org/10.1021/acs.accounts.6b00435.

[3] G. van Meer, D.R. Voelker, G.W. Feigenson, Membrane lipids: where they are and how they behave, Nat. Rev. Mol. Cell Biol. 9 (2008) 112–124. https://doi.org/10.1038/nrm2330.

[4] A. Laganowsky, E. Reading, T.M. Allison, M.B. Ulmschneider, M.T. Degiacomi, A.J. Baldwin, C.V. Robinson, Membrane proteins bind lipids selectively to modulate their structure and function, Nature. 510 (2014) 172–175. https://doi.org/10.1038/nature13419.

[5] V. Corradi, B.I. Sejdiu, H. Mesa-Galloso, H. Abdizadeh, S.Yu. Noskov, S.J. Marrink, D.P. Tieleman, Emerging Diversity in Lipid–Protein Interactions, Chem. Rev. 119 (2019) 5775–5848. https://doi.org/10.1021/acs.chemrev.8b00451.

[6] F.-X. Contreras, A.M. Ernst, F. Wieland, B. Brügger, Specificity of Intramembrane Protein–Lipid Interactions, Cold Spring Harb. Perspect. Biol. 3 (2011) a004705. https://doi.org/10.1101/cshperspect.a004705.

[7] C.W. Cotman, M.L. Blank, A. Moehl, F. Snyder, Lipid composition of synaptic plasma membranes isolated from rat brain by zonal centrifugation, Biochemistry. 8 (1969) 4606–4612. https://doi.org/10.1021/bi00839a056.

[8] A. Yamashita, Y. Hayashi, Y. Nemoto-Sasaki, M. Ito, S. Oka, T. Tanikawa, K. Waku, T. Sugiura, Acyltransferases and transacylases that determine the fatty acid composition of glycerolipids and the metabolism of bioactive lipid mediators in mammalian cells and model organisms, Prog. Lipid Res. 53 (2014) 18–81. https://doi.org/10.1016/j.plipres.2013.10.001.

[9] S.J. Marrink, V. Corradi, P.C.T. Souza, H.I. Ingólfsson, D.P. Tieleman, M.S.P. Sansom, Computational Modeling of Realistic Cell Membranes, Chem. Rev. 119 (2019) 6184–6226. https://doi.org/10.1021/acs.chemrev.8b00460.

[10] H.I. Ingólfsson, M.N. Melo, F.J. van Eerden, C. Arnarez, C.A. Lopez, T.A. Wassenaar, X. Periole, A.H. de Vries, D.P. Tieleman, S.J. Marrink, Lipid Organization of the Plasma Membrane, J. Am. Chem. Soc. 136 (2014) 14554–14559. https://doi.org/10.1021/ja507832e.

[11] H.I. Ingólfsson, T.S. Carpenter, H. Bhatia, P.-T. Bremer, S.J. Marrink, F.C. Lightstone, Computational Lipidomics of the Neuronal Plasma Membrane, Biophys. J. 113 (2017) 2271–2280. https://doi.org/10.1016/j.bpj.2017.10.017.

[12] V. Corradi, E. Mendez-Villuendas, H.I. Ingólfsson, R.-X. Gu, I. Siuda, M.N. Melo, A. Moussatova, L.J. DeGagné, B.I. Sejdiu, G. Singh, T.A. Wassenaar, K. Delgado Magnero, S.J. Marrink, D.P. Tieleman, Lipid–Protein Interactions Are Unique Fingerprints for Membrane Proteins, ACS Cent. Sci. 4 (2018) 709–717. https://doi.org/10.1021/acscentsci.8b00143.

[13] K.A. Wilson, H.I. MacDermott-Opeskin, E. Riley, Y. Lin, M.L. O’Mara, Understanding the Link between Lipid Diversity and the Biophysical Properties of the Neuronal Plasma Membrane, Biochemistry. 59 (2020) 3010–3018. https://doi.org/10.1021/acs.biochem.0c00524.

[14] P.A. Bala, J. Foster, L. Carvelli, L.K. Henry, SLC6 Transporters: Structure, Function, Regulation, Disease Association and Therapeutics, Mol. Aspects Med. 34 (2013) 197–219. https://doi.org/10.1016/j.mam.2012.07.002.

[15] M.J. Abraham, T. Murtola, R. Schulz, S. Páll, J.C. Smith, B. Hess, E. Lindahl, GROMACS: High performance molecular simulations through multi-level parallelism from laptops to supercomputers, SoftwareX. 1–2 (2015) 19–25. https://doi.org/10.1016/j.softx.2015.06.001.

[16] D.H. de Jong, G. Singh, W.F.D. Bennett, C. Arnarez, T.A. Wassenaar, L.V. Schäfer, X. Periole, D.P. Tieleman, S.J. Marrink, Improved Parameters for the Martini Coarse-Grained Protein Force Field, J. Chem. Theory Comput. 9 (2013) 687–697. https://doi.org/10.1021/ct300646g.

[17] K.H. Wang, A. Penmatsa, E. Gouaux, Neurotransmitter and psychostimulant recognition by the dopamine transporter, Nature. 521 (2015) 322–327. https://doi.org/10.1038/nature14431.

[18] J.A. Coleman, E.M. Green, E. Gouaux, X-ray structures and mechanism of the human serotonin transporter, Nature. 532 (2016) 334–339. https://doi.org/10.1038/nature17629.

[19] N. Subramanian, A.J. Scopelitti, J.E. Carland, R.M. Ryan, M.L. O’Mara, R.J. Vandenberg, Identification of a 3rd Na+ Binding Site of the Glycine Transporter, GlyT2, PLOS ONE. 11 (2016) e0157583. https://doi.org/10.1371/journal.pone.0157583.

[20] S.J. Marrink, H.J. Risselada, S. Yefimov, D.P. Tieleman, A.H. de Vries, The MARTINI Force Field: Coarse Grained Model for Biomolecular Simulations, J. Phys. Chem. B. 111 (2007) 7812–7824. https://doi.org/10.1021/jp071097f.

[21] T.A. Wassenaar, H.I. Ingólfsson, R.A. Böckmann, D.P. Tieleman, S.J. Marrink, Computational Lipidomics with insane: A Versatile Tool for Generating Custom Membranes for Molecular Simulations, J. Chem. Theory Comput. 11 (2015) 2144–2155. https://doi.org/10.1021/acs.jctc.5b00209.

[22] H.I. Ingólfsson, T.S. Carpenter, H. Bhatia, P.-T. Bremer, S.J. Marrink, F.C. Lightstone, Computational Lipidomics of the Neuronal Plasma Membrane, Biophys. J. 113 (2017) 2271–2280. https://doi.org/10.1016/j.bpj.2017.10.017.

[23] D.H. de Jong, S. Baoukina, H.I. Ingólfsson, S.J. Marrink, Martini straight: Boosting performance using a shorter cutoff and GPUs, Comput. Phys. Commun. 199 (2016) 1–7. https://doi.org/10.1016/j.cpc.2015.09.014.

[24] G. Bussi, D. Donadio, M. Parrinello, Canonical sampling through velocity-rescaling, J. Chem. Phys. 126 (2007) 014101. https://doi.org/10.1063/1.2408420.

[25] M. Parrinello, A. Rahman, Polymorphic transitions in single crystals: A new molecular dynamics method, J. Appl. Phys. 52 (1981) 7182–7190. https://doi.org/10.1063/1.328693.

[26] N. Castillo, L. Monticelli, J. Barnoud, D.P. Tieleman, Free energy of WALP23 dimer association in DMPC, DPPC, and DOPC bilayers, Chem. Phys. Lipids. 169 (2013) 95–105. https://doi.org/10.1016/j.chemphyslip.2013.02.001.

[27] E.J. Maginn, R.A. Messerly, D.J. Carlson, D.R. Roe, J.R. Elliott, Best Practices for Computing Transport Properties 1. Self-Diffusivity and Viscosity from Equilibrium Molecular Dynamics [Article v1.0], Living J. Comput. Mol. Sci. 1 (2018) 6324. https://doi.org/10.33011/livecoms.1.1.6324.

[28] W. Humphrey, A. Dalke, K. Schulten, VMD: Visual molecular dynamics, J. Mol. Graph. 14 (1996) 33–38. https://doi.org/10.1016/0263-7855(96)00018-5.

[29] N. Michaud-Agrawal, E.J. Denning, T.B. Woolf, O. Beckstein, MDAnalysis: A toolkit for the analysis of molecular dynamics simulations, J. Comput. Chem. 32 (2011) 2319–2327. https://doi.org/10.1002/jcc.21787.

[30] R.J. Gowers, M. Linke, J. Barnoud, T.J.E. Reddy, M.N. Melo, S.L. Seyler, J. Domański, D.L. Dotson, S. Buchoux, I.M. Kenney, O. Beckstein, MDAnalysis: A Python Package for the Rapid Analysis of Molecular Dynamics Simulations, Proc. 15th Python Sci. Conf. (2016) 98–105. https://doi.org/10.25080/Majora-629e541a-00e.

[31] S.N. Mostyn, J.E. Carland, S. Shimmon, R.M. Ryan, T. Rawling, R.J. Vandenberg, Synthesis and Characterization of Novel Acyl-Glycine Inhibitors of GlyT2, ACS Chem. Neurosci. 8 (2017) 1949–1959. https://doi.org/10.1021/acschemneuro.7b00105.

[32] M. L’hirondel, A. Chéramy, G. Godeheu, J. Glowinski, Effects of Arachidonic Acid on Dopamine Synthesis, Spontaneous Release, and Uptake in Striatal Synaptosomes from the Rat, J. Neurochem. 64 (1995) 1406–1409. https://doi.org/10.1046/j.1471-4159.1995.64031406.x.

[33] Regulation of the functional activity of the human dopamine transporter by the arachidonic acid pathway, Eur. J. Pharmacol. 315 (1996) 345–354. https://doi.org/10.1016/S0014-2999(96)00646-2.

[34] R.M. Vernon, P.A. Chong, B. Tsang, T.H. Kim, A. Bah, P. Farber, H. Lin, J.D. Forman-Kay, Pi-Pi contacts are an overlooked protein feature relevant to phase separation, ELife. 7 (2018) e31486. https://doi.org/10.7554/eLife.31486.

[35] K.A. Wilson, J.L. Kellie, S.D. Wetmore, DNA–protein π-interactions in nature: abundance, structure, composition and strength of contacts between aromatic amino acids and DNA nucleobases or deoxyribose sugar, Nucleic Acids Res. 42 (2014) 6726–6741. https://doi.org/10.1093/nar/gku269.

[36] K.A. Wilson, S.D. Wetmore, Combining crystallographic and quantum chemical data to understand DNA-protein π-interactions in nature, Struct. Chem. 28 (2017) 1487–1500. https://doi.org/10.1007/s11224-017-0954-7.

[37] K.L. Hudson, G.J. Bartlett, R.C. Diehl, J. Agirre, T. Gallagher, L.L. Kiessling, D.N. Woolfson, Carbohydrate–Aromatic Interactions in Proteins, J. Am. Chem. Soc. 137 (2015) 15152–15160. https://doi.org/10.1021/jacs.5b08424.

[38] L. Montalvillo-Jiménez, A.G. Santana, F. Corzana, G. Jiménez-Osés, J. Jiménez-Barbero, A.M. Gómez, J.L. Asensio, Impact of Aromatic Stacking on Glycoside Reactivity: Balancing CH/π and Cation/π Interactions for the Stabilization of Glycosyl-Oxocarbenium Ions, J. Am. Chem. Soc. 141 (2019) 13372–13384. https://doi.org/10.1021/jacs.9b03285.

[39] K.M. Sanchez, G. Kang, B. Wu, J.E. Kim, Tryptophan-Lipid Interactions in Membrane Protein Folding Probed by Ultraviolet Resonance Raman and Fluorescence Spectroscopy, Biophys. J. 100 (2011) 2121–2130. https://doi.org/10.1016/j.bpj.2011.03.018.

[40] A.N. Belovich, J.I. Aguilar, S.J. Mabry, M.H. Cheng, D. Zanella, P.J. Hamilton, D.J. Stanislowski, A. Shekar, J.D. Foster, I. Bahar, H.J.G. Matthies, A. Galli, A network of phosphatidylinositol (4,5)-bisphosphate (PIP 2) binding sites on the dopamine transporter regulates amphetamine behavior in Drosophila Melanogaster, Mol. Psychiatry. (2019) 1–14. https://doi.org/10.1038/s41380-019-0620-0.

[41] W.C. Hong, S.G. Amara, Membrane cholesterol modulates the outward facing conformation of the dopamine transporter and alters cocaine binding, J. Biol. Chem. 285 (2010) 32616–32626. https://doi.org/10.1074/jbc.M110.150565.

[42] K.T. Jones, J. Zhen, M.E.A. Reith, Importance of cholesterol in dopamine transporter function, J. Neurochem. 123 (2012) 700–715. https://doi.org/10.1111/jnc.12007.

[43] H. Bjerregaard, K. Severinsen, S. Said, O. Wiborg, S. Sinning, A Dualistic Conformational Response to Substrate Binding in the Human Serotonin Transporter Reveals a High Affinity State for Serotonin, J. Biol. Chem. 290 (2015) 7747–7755. https://doi.org/10.1074/jbc.M114.573477.

[44] S.M. Scanlon, D.C. Williams, P. Schloss, Membrane Cholesterol Modulates Serotonin Transporter Activity, Biochemistry. 40 (2001) 10507–10513. https://doi.org/10.1021/bi010730z.

[45] C.B. Divito, S.G. Amara, Close Encounters of the Oily Kind: Regulation of Transporters by Lipids, Mol. Interv. 9 (2009) 252–262. https://doi.org/10.1124/mi.9.5.8.

[46] A. Penmatsa, K.H. Wang, E. Gouaux, X-ray structure of dopamine transporter elucidates antidepressant mechanism, Nature. 503 (2013) 85–90. https://doi.org/10.1038/nature12533.

[47] A. Penmatsa, K.H. Wang, E. Gouaux, X-ray structures of Drosophila dopamine transporter in complex with nisoxetine and reboxetine, Nat. Struct. Mol. Biol. 22 (2015) 506–508. https://doi.org/10.1038/nsmb.3029.

[48] T. Zeppelin, L.K. Ladefoged, S. Sinning, X. Periole, B. Schiøtt, A direct interaction of cholesterol with the dopamine transporter prevents its out-to-inward transition, PLOS Comput. Biol. 14 (2018) e1005907. https://doi.org/10.1371/journal.pcbi.1005907.

[49] J. Fantini, F.J. Barrantes, How cholesterol interacts with membrane proteins: an exploration of cholesterol-binding sites including CRAC, CARC, and tilted domains, Front. Physiol. 4 (2013). https://doi.org/10.3389/fphys.2013.00031.

[50] J. Fantini, D. Carlus, N. Yahi, The fusogenic tilted peptide (67–78) of α-synuclein is a cholesterol binding domain, Biochim. Biophys. Acta BBA - Biomembr. 1808 (2011) 2343–2351. https://doi.org/10.1016/j.bbamem.2011.06.017.

[51] M. Nishio, Y. Umezawa, M. Hirota, Y. Takeuchi, The CH/π interaction: Significance in molecular recognition, Tetrahedron. 51 (1995) 8665–8701. https://doi.org/10.1016/0040-4020(94)01066-9.

[52] S. Park, A.H. Beaven, J.B. Klauda, W. Im, How Tolerant are Membrane Simulations with Mismatch in Area per Lipid between Leaflets?, J. Chem. Theory Comput. 11 (2015) 3466–3477. https://doi.org/10.1021/acs.jctc.5b00232.

[53] J.C. Mathai, S. Tristram-Nagle, J.F. Nagle, M.L. Zeidel, Structural Determinants of Water Permeability through the Lipid Membrane, J. Gen. Physiol. 131 (2008) 69–76. https://doi.org/10.1085/jgp.200709848.

[54] I. Ermilova, A.P. Lyubartsev, Extension of the Slipids Force Field to Polyunsaturated Lipids, J. Phys. Chem. B. 120 (2016) 12826–12842. https://doi.org/10.1021/acs.jpcb.6b05422.

[55] A. Prinetti, N. Loberto, V. Chigorno, S. Sonnino, Glycosphingolipid behaviour in complex membranes, Biochim. Biophys. Acta BBA - Biomembr. 1788 (2009) 184–193. https://doi.org/10.1016/j.bbamem.2008.09.001.

[56] W.F.D. Bennett, J.L. MacCallum, M.J. Hinner, S.J. Marrink, D.P. Tieleman, Molecular View of Cholesterol Flip-Flop and Chemical Potential in Different Membrane Environments, J. Am. Chem. Soc. 131 (2009) 12714–12720. https://doi.org/10.1021/ja903529f.

[57] F. de Meyer, B. Smit, Effect of cholesterol on the structure of a phospholipid bilayer, Proc. Natl. Acad. Sci. U. S. A. 106 (2009) 3654–3658. https://doi.org/10.1073/pnas.0809959106.

[58] A. Leftin, T.R. Molugu, C. Job, K. Beyer, M.F. Brown, Area per Lipid and Cholesterol Interactions in Membranes from Separated Local-Field 13C NMR Spectroscopy, Biophys. J. 107 (2014) 2274–2286. https://doi.org/10.1016/j.bpj.2014.07.044.

[59] J.M. Smaby, M.M. Momsen, H.L. Brockman, R.E. Brown, Phosphatidylcholine acyl unsaturation modulates the decrease in interfacial elasticity induced by cholesterol., Biophys. J. 73 (1997) 1492–1505.

