## Supporting Information Figures 1-10, expanded methods for "Lipid fingerprints are similar between SLC6 transporters in the neuronal membrane"

**Table of Contents**

Expanded methods ........................................ S2

Leaflet detection ........................................ S2

Lipid flip-flop ........................................ S3

Area per lipid ........................................ S3

Depletion-enrichment index ........................................ S3

References ........................................ S4

Figure S1. Membrane composition by saturation ........................................ S6

Figure S2. The mean enrichment-depletion index of lipids in the protein annulus ........................................ S6

Figure S3. Areas around the protein with at least 0.005 Å$^{-3}$ density of PE lipids ........................................ S7

Figure S4. The natural abundance of each amino acid in each protein ........................................ S7

Figure S5. Areas around the protein with at least 0.005 Å$^{-3}$ density of GS lipids ........................................ S7

Figure S6. Areas around the protein with at least 0.005 Å$^{-3}$ density of PI lipids ........................................ S8

Figure S8. The mean area per lipid of lipids in the protein annulus grouped by tail saturation ........................................ S9

Figure S9. Membrane thickness across the x-y plane of the model POPC-CHOL membraneS9

Figure S10. The spectral-clustering leaflet detection algorithm ........................................ S10

**Expanded methods**

All code below is built on the *MDAnalysis* [1,2] package. It is available at https://github.com/OMaraLab/SLC6_lipid_fingerprints. Images were created using VMD and graphs were plotted using *seaborn* [3] and *matplotlib* [4].

***Leaflet detection***

Determining lipid membership within leaflets was crucial to determining the flip-flop rate, area per lipid, and lipid depletion-enrichment indices. We applied spectral clustering [5–7] using the *scikit-learn* [8] package to an affinity matrix that incorporated both the distance between lipid headgroups, and the similarity of each lipid orientation as quantified by the cosine similarity of the lipid orientations.

We first defined two points for each lipid: the center-of-geometry of the selected headgroup/s, and the center-of-geometry of the remaining particles (Figure S10a-b). The orientation of each lipid was approximated as the vector between these two points, originating from the headgroup center (Figure S10b). A cosine similarity was calculated between each lipid orientation vector (Figure S10c). We clipped the cosine similarity to a range of [-0.6, 0.6] to avoid overly weighting extremely parallel or anti-parallel lipids; we then normalised this to a range of [0, 1] such that two orthogonal vectors have a similarity of 0.5, antiparallel vectors have a similarity of 0, and parallel vectors are represented by 1.

An original distance matrix was calculated between the centers-of-geometry of the selected headgroup of each lipid within a cut-off radius of 60 Å (Figure S10d). We then constructed another distance matrix consisting of the original distances projected onto the vector of the lipid orientation (Figure S10d). This second matrix reduces the distance separation of lipids on the plane of (“next to”) the central lipid, but preserves the separation of orthogonal (“below or above”) lipids. This projected distance matrix was not symmetric, due to the different orientations of the two lipids in a pair. It was symmetrised by taking the average of the upper and lower diagonal values. The final distance matrix is the sum of the original distances and the projected distances. As the distance values between lipid pairs that are further than the cut-off radius are still undefined in this final matrix, they are set to double the cut-off radius as the placeholder value. The distance matrix was transformed into a suitable similarity matrix using a Gaussian kernel with a delta value of 20. The final affinity matrix was constructed as the product of the distance similarity matrix and the cosine similarity matrix.

This new method of leaflet detection was necessary to accurately cluster lipids into two leaflets, especially around the protein. Other leaflet detection methods, such as the graph-construction method of Michaud-Agrawal *et al.*, 2011 [1], do not account for lipid orientation; membrane features such as the space occupied by a transmembrane protein, and cholesterol molecules lying near the inter-leaflet interface, can cause poor performance and the failure to assign all lipids to two dominant leaflets (Figure S10e). Methods such as the FATSLiM algorithm [9], which constructs a local normal for each lipid from the nearest neighbors, can be confounded by the variable orientations and positioning of cholesterol lipids; cut-off distances that allow local normals to be constructed from lipids in both leaflets can result in normal vectors that lie in the plane of the bilayer, rather than being orthogonal. The spectral clustering algorithm incorporates both distances and lipid orientations, and moreover increases the separation between the two leaflets by adding the projected distances between lipids.

This allows leaflets to be readily detected even when cholesterol molecules lie near the inter-leaflet interface and when a protein is positioned in the membrane (Figure S10f).

For all methods requiring leaflet-wise analysis, lipids were first separated into leaflets. If multiple headgroups were specified for each lipid, we used the center-of-geometry of the groups as the lipid headgroup coordinate. Unless otherwise stated, we specified headgroups to be the ROH, PO4, GL1, GL2, AM1, AM2, NC3, GM1–9, C3 beads for all lipids, in addition to the C1 and C2 beads for the GS lipids.

***Lipid flip-flop***

Changes in orientation and being located near the inter-leaflet interface can cause a lipid to rapidly change membership between leaflets as detected by the spectral clustering method, so we defined an interstitial space of 16 Å between the leaflets akin to the method in Ingólfsson *et al.*, 2017 [10] and Wilson *et al.*, 2020 [11]. Transitions from a leaflet into the interstitial space were not considered to be translocations. Lipids were first separated into leaflets using the spectral clustering method described above. Centers-of-geometry were calculated for each of these leaflet groups. An inter-leaflet center was calculated as the midpoint of the leaflet centers. The interstitial space was defined to lie within 8 Å from the inter-leaflet center. The lipid $L_i$ was then re-assigned to either its initial leaflet, as determined via spectral clustering, or the interstitial space. Changes in leaflet membership were deemed to be a translocation (a "flip" or a "flop", depending on direction). The flip-flop rate was calculated to be the total number of translocations, over the number of cholesterol molecules in the system, and the time-frame considered. Rates were reported in numbers of translocations per $10^6$ s.

***Area per lipid***

Lipids were first separated via the spectral clustering method described above. We then restricted the headgroups considered for the analysis to the PO4, GL1, ROH, and AM1 beads. All neighbouring lipid headgroups within the given cut-off (40 Å) of each lipid were assumed to be planar to the central lipid. We thereby calculated the plane of best fit for each lipid and projected the coordinates of the neighbouring lipids and neighbouring protein atoms onto the plane. Voronoi diagrams were constructed from these points using *scipy.spatial.Voronoi,* which wraps the Qhull library [12]. The area per lipid was calculated as the area of the corresponding cell. The area per lipid was calculated for each lipid molecule separately; values were aggregated by lipid species, headgroup, and tail saturation to draw conclusions. Cholesterol molecules were included in the analysis to account for effects on other lipid molecules, but values were not reported due to its propensity to lie near the inter-leaflet interface and flip-flop.

***Depletion-enrichment index***

The lipid depletion-enrichment index (DEI) metric was drawn from Corradi *et al.*, 2018 [13]. All DEIs were calculated per-leaflet, to account for flip-flop behaviour. Lipids were assigned to each leaflet following the spectral clustering method described above. We then restricted the beads per lipid used in the analysis to the PO4, ROH, GL1, GL2, AM1, AM2, C3, and NC3 beads for computational efficiency. For each lipid type $L$ we used a binomial distribution to approximate the likelihood of finding $k$ number of $L$ lipids, in the group of $n$ lipids within a cut-off radius around the protein, given a total $K$ lipids of type $L$ in a leaflet with $N$ lipids of any type. Distances were calculated only to the beads specified above, rather to every bead of each lipid; however, every bead of the protein was considered.

$$\Pr(X_L = k) = \binom{n}{k} p^k (1-p)^{n-k}$$

We fit the aggregate of these events over a trajectory to the convolution of binomial distributions for each frame. The convolution of binomial distributions is a new binomial distribution. The probability distribution of the DEI is then the ratio of the observed binomial distribution with $p_{obs}$, divided by the binomial distribution of null hypothesis $p_0$. This ratio is approximated as a log-normal distribution, where the DEI is the median value:

$$DEI = \frac{p_{obs}}{p_0}$$

$$p_0 = \frac{K_L}{N}$$

We found that using a hard cut-off to analyse the lipid enrichment and depletion profiles around a protein resulted in extreme initial index values. Ideally these converge to representative values as sampling increases, but the analysis is nonetheless unduly affected by increasing time intervals between the frames analysed. In order to capture and appropriately weight near-contact events that may represent contacts during the simulation that lie between the frames analysed, we added a soft cut-off buffer zone. While contacts within the hard cut-off are treated as successful contacts with a weighting of 1; lipids within the buffer zone are weighted according to a Gaussian function that reduces to 0 at the outer edge of the buffer zone. All values reported here were calculated with a hard cut-off of 6 Å and a buffer zone of 2 Å. As the DEI reports the behaviour of a group of lipids, the DEI values were separately calculated to describe the enrichment of different lipid species, headgroups, and tail saturations.

***References***


[1] N. Michaud-Agrawal, E.J. Denning, T.B. Woolf, O. Beckstein, MDAnalysis: A toolkit for the analysis of molecular dynamics simulations, J. Comput. Chem. 32 (2011) 2319–2327. https://doi.org/10.1002/jcc.21787.

[2] R.J. Gowers, M. Linke, J. Barnoud, T.J.E. Reddy, M.N. Melo, S.L. Seyler, J. Domański, D.L. Dotson, S. Buchoux, I.M. Kenney, O. Beckstein, MDAnalysis: A Python Package for the Rapid Analysis of Molecular Dynamics Simulations, Proceedings of the 15th Python in Science Conference. (2016) 98–105. https://doi.org/10.25080/Majora-629e541a-00e.

[3] M. Waskom, the seaborn development team, mwaskom/seaborn, Zenodo, 2020. https://doi.org/10.5281/zenodo.592845.

[4] J.D. Hunter, Matplotlib: A 2D graphics environment, Computing in Science & Engineering. 9 (2007) 90–95. https://doi.org/10.1109/MCSE.2007.55.

[5] J. Shi, J. Malik, Normalized cuts and image segmentation, 2000.

[6] U.V. Luxburg, A Tutorial on Spectral Clustering, 2007.

[7] Yu, Shi, Multiclass spectral clustering, in: Proceedings Ninth IEEE International Conference on Computer Vision, 2003: pp. 313–319 vol.1. https://doi.org/10.1109/ICCV.2003.1238361.

[8] F. Pedregosa, G. Varoquaux, A. Gramfort, V. Michel, B. Thirion, O. Grisel, M. Blondel, P. Prettenhofer, R. Weiss, V. Dubourg, J. Vanderplas, A. Passos, D. Cournapeau, M. Brucher, M. Perrot, E. Duchesnay, Scikit-learn: Machine Learning in Python, Journal of Machine Learning Research. 12 (2011) 2825–2830.

[9] S. Buchoux, FATSLiM: a fast and robust software to analyze MD simulations of membranes, Bioinformatics. 33 (2017) 133–134. https://doi.org/10.1093/bioinformatics/btw563.

[10] H.I. Ingólfsson, T.S. Carpenter, H. Bhatia, P.-T. Bremer, S.J. Marrink, F.C. Lightstone, Computational Lipidomics of the Neuronal Plasma Membrane, Biophysical Journal. 113 (2017) 2271–2280. https://doi.org/10.1016/j.bpj.2017.10.017.

[11] K.A. Wilson, H.I. MacDermott-Opeskin, E. Riley, Y. Lin, M.L. O'Mara, Understanding the Link between Lipid Diversity and the Biophysical Properties of the Neuronal Plasma Membrane, Biochemistry. 59 (2020) 3010–3018. https://doi.org/10.1021/acs.biochem.0c00524.
[12] C.B. Barber, D.P. Dobkin, H. Huhdanpaa, The Quickhull algorithm for convex hulls, Acm Transactions on Mathematical Software. 22 (1996) 469–483.
[13] V. Corradi, E. Mendez-Villuendas, H.I. Ingólfsson, R.-X. Gu, I. Siuda, M.N. Melo, A. Moussatova, L.J. DeGagné, B.I. Sejdiu, G. Singh, T.A. Wassenaar, K. Delgado Magnero, S.J. Marrink, D.P. Tieleman, Lipid–Protein Interactions Are Unique Fingerprints for Membrane Proteins, ACS Cent Sci. 4 (2018) 709–717. https://doi.org/10.1021/acscentsci.8b00143.

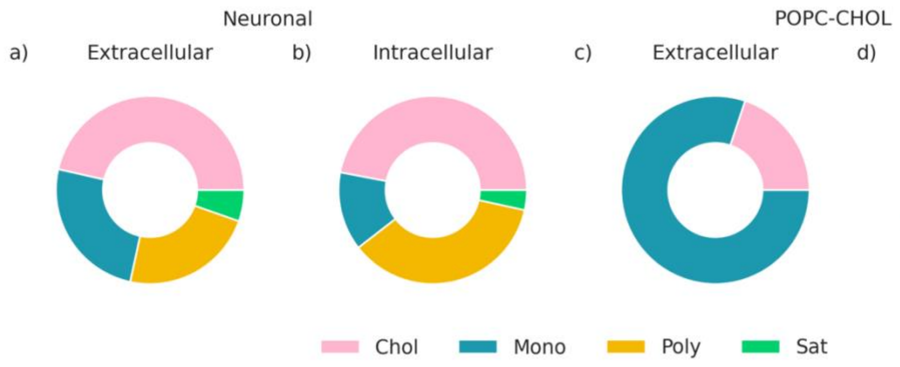


**Figure S1**. Membrane composition by saturation of **a)** the extracellular leaflet of the neuronal membrane; **b)** the intracellular leaflet of the neuronal membrane; **c)** the extracellular leaflet of the model POPC-CHOL membrane; and **d)** the intracellular leaflet of the model POPC-CHOL membrane. Full composition and definition of acronyms are shown in Table S1.

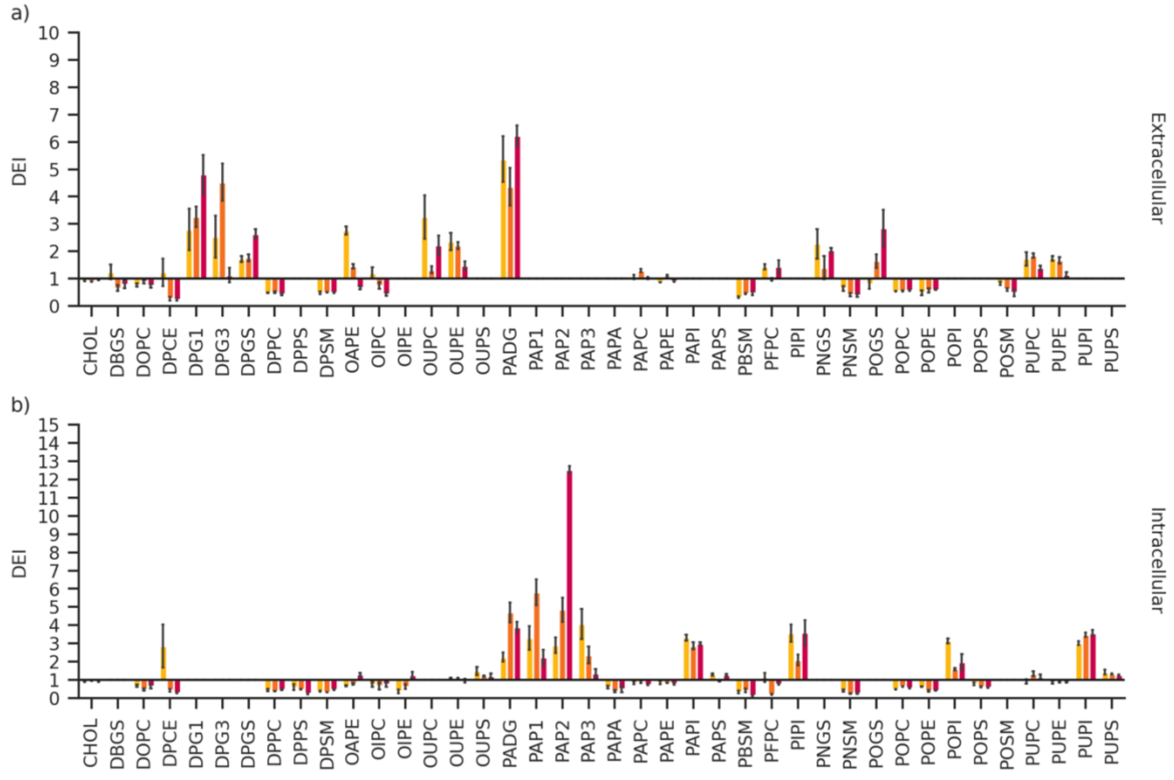


**Figure S2.** The mean enrichment-depletion index of lipids in the protein annulus in the **a)** extracellular leaflet; and **b)** the intracellular leaflet. The standard error from the mean is represented by the error bars. DEI values of each lipid species are shown in Table S2. Yellow: dDAT. Orange: GlyT2. Red: hSERT.

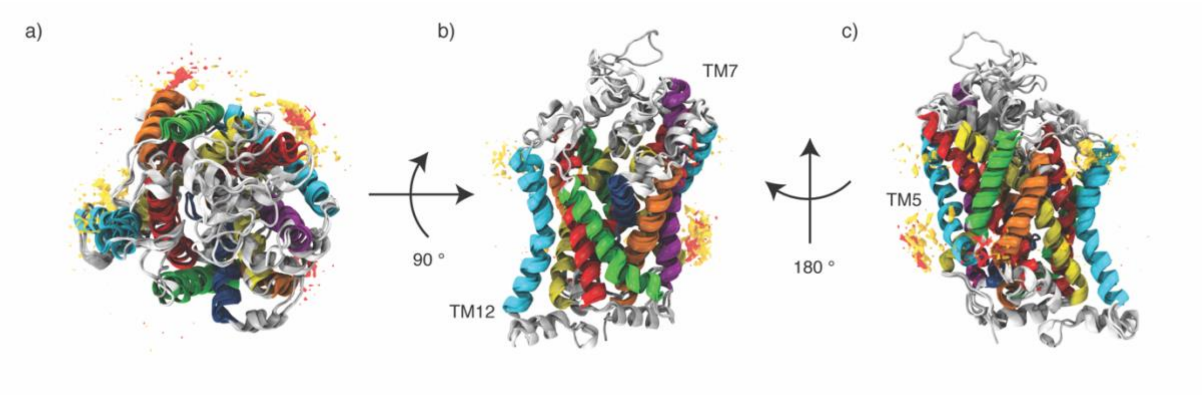


**Figure S3.** Areas around the protein with at least 0.005 Å$^{-3}$ density of PE lipids. dDAT, GlyT2, and hSERT are superimposed on each other. Yellow: areas of high density around dDAT. Orange: areas of high density around GlyT2. Red: areas of high density around hSERT. **a)** top-down view **b)** side view **c)** rotated side view.

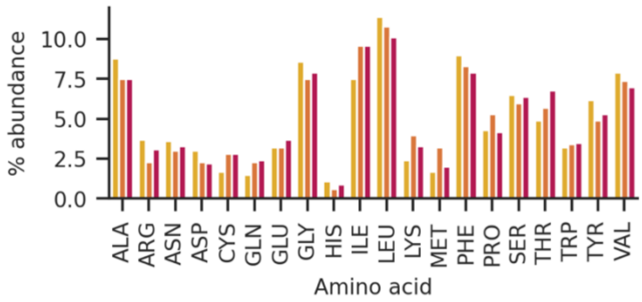


**Figure S4.** The natural abundance of each amino acid in each protein. Yellow: dDAT. Orange: GlyT2. Red: hSERT.

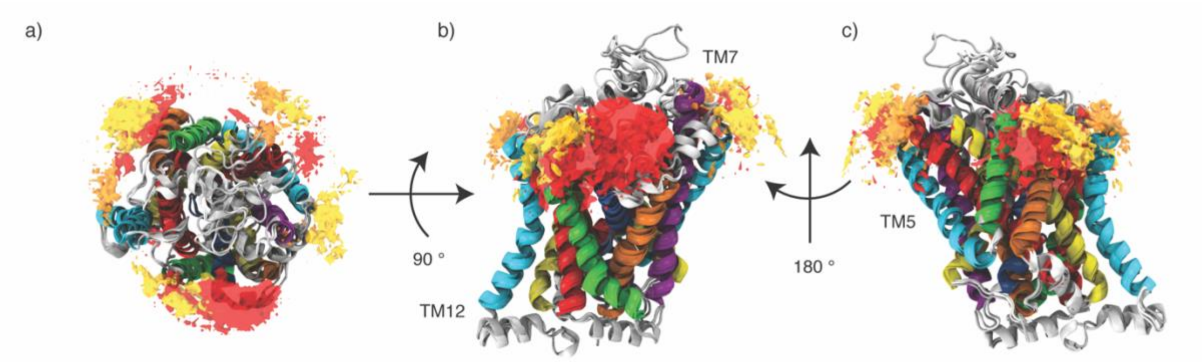


**Figure S5.** Areas around the protein with at least 0.005 Å$^{-3}$ density of GS lipids. dDAT, GlyT2, and hSERT are superimposed on each other. Yellow: areas of high density around dDAT. Orange: areas of high density around GlyT2. Red: areas of high density around hSERT. **a)** top-down view **b)** side view **c)** rotated side view.

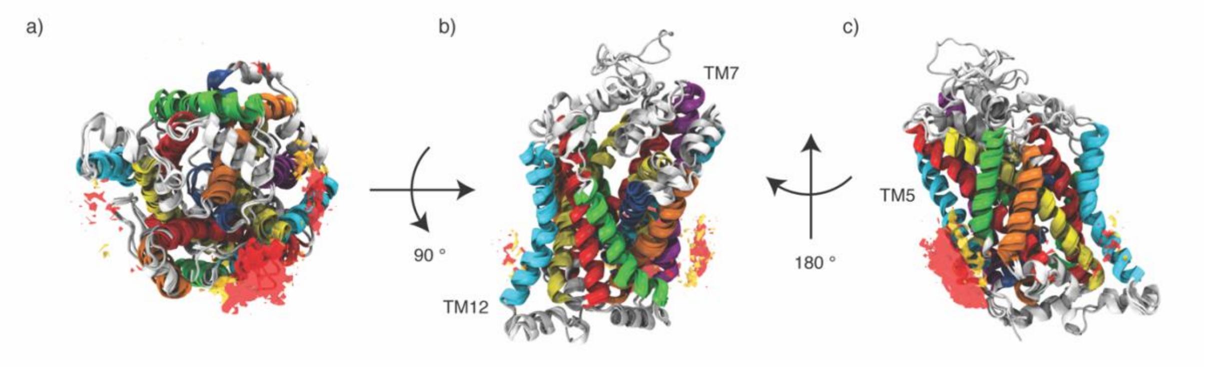


**Figure S6**. Areas around the protein with at least 0.005 Å$^{-3}$ density of PI lipids. dDAT, GlyT2, and hSERT are superimposed on each other. Yellow: areas of high density around dDAT. Orange: areas of high density around GlyT2. Red: areas of high density around hSERT. **a)** bottom-up view **b)** side view **c)** rotated side view.

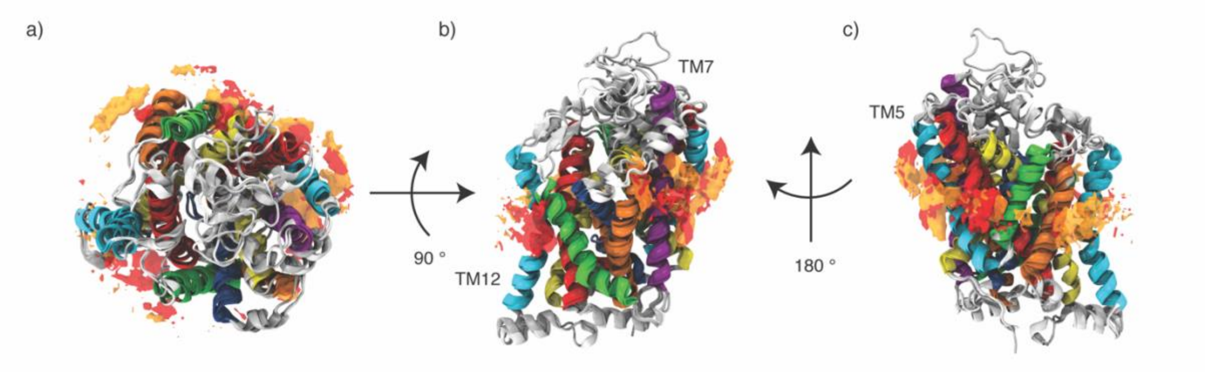


**Figure S7**. Areas around the protein with at least 0.010 Å-3 density of cholesterol. dDAT, GlyT2, and hSERT are superimposed on each other. Yellow: areas of high density around dDAT. Orange: areas of high density around GlyT2. Red: areas of high density around hSERT. **a)** top-down view **b)** side view **c)** rotated side view.

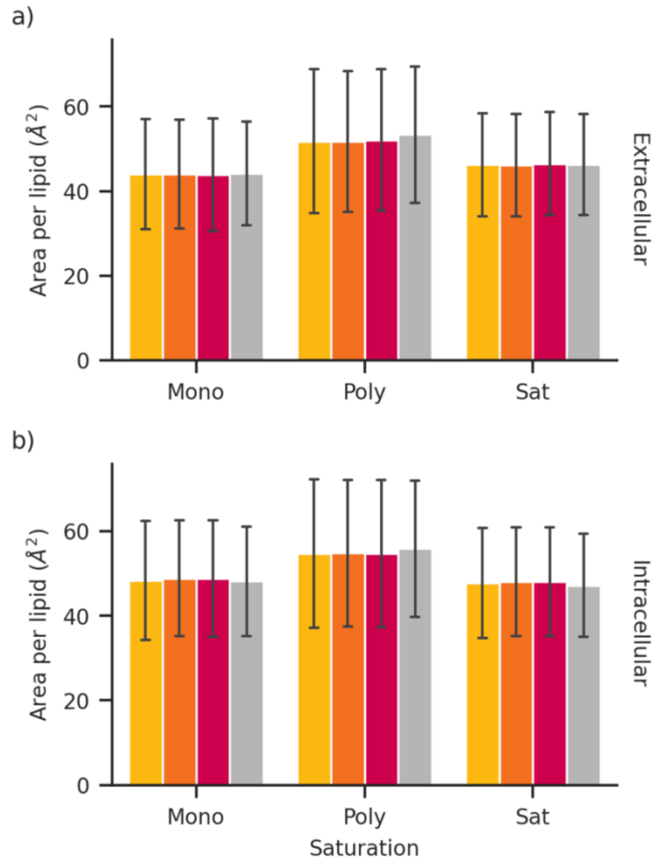


**Figure S8**. The mean area per lipid of lipids in the protein annulus grouped by tail saturation, in the **a)** extracellular leaflet; and **b)** the intracellular leaflet. The standard deviation is represented by the error bars. Yellow: dDAT. Orange: GlyT2. Red: hSERT. None: no protein [11].

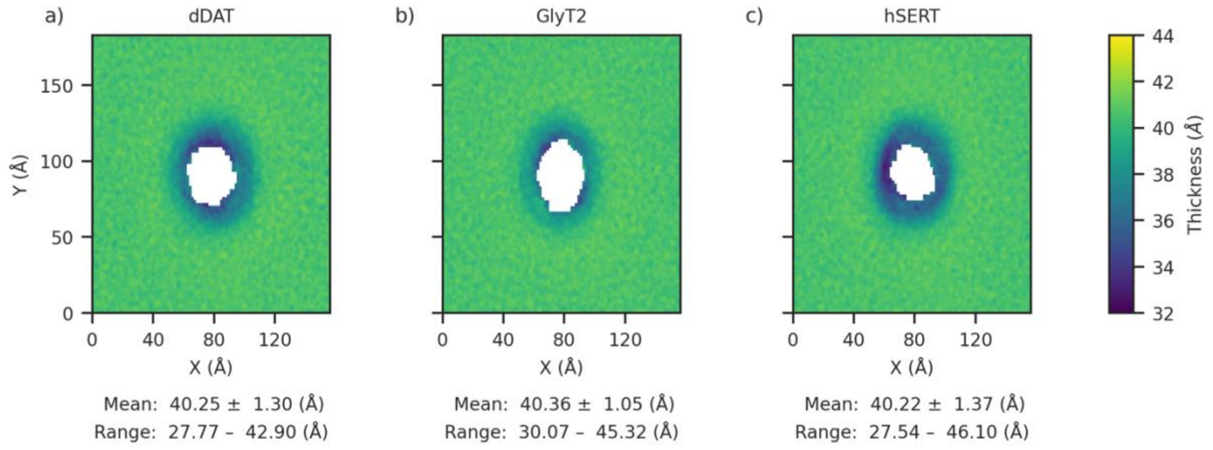


**Figure S9.** Membrane thickness across the x-y plane of the model POPC-CHOL membrane embedded with a) dDAT; b) GlyT2; and c) hSERT. Proteins are not aligned to each other. The colour bar is truncated and does not represent the full range of values. Average and standard deviation of the thickness is shown below each plot, followed by the minimum and maximum values.

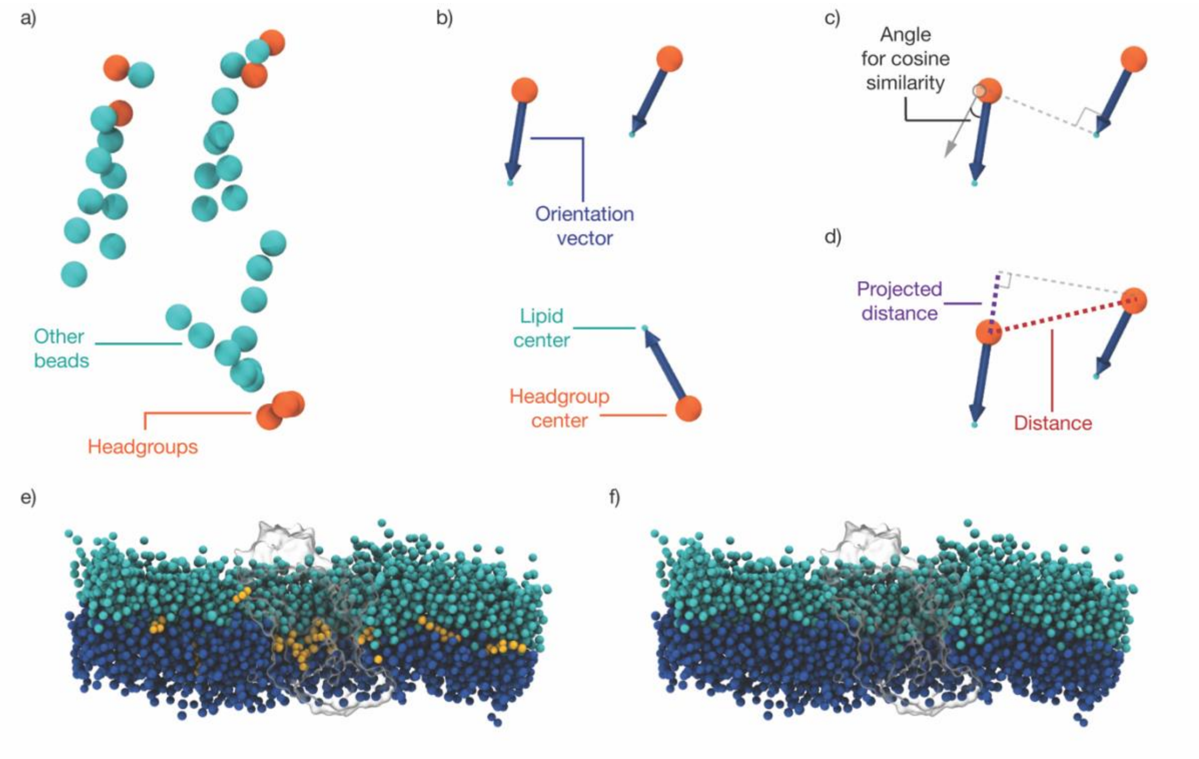


**Figure S10.** The spectral-clustering leaflet detection algorithm. **a)** Headgroups (orange) are defined for each lipid. **b)** The center-of-geometry is calculated for headgroups (orange) and the remaining beads (cyan). A vector describing the orientation of the lipid (dark blue) is defined from the headgroup center to the lipid center. **c)** The cosine similarity between each orientation vector (dark blue) is calculated. **d)** The actual distance between each headgroup center is calculated (red). The projected distance between each headgroup, onto each lipid orientation vector, is calculated (purple). **e)** Performance of the graph-construction leaflet-detection algorithm in Michaud-Agrawal *et al.*, 2011 [1] on the neuronal membrane. Cyan: extracellular leaflet. Dark blue: intracellular leaflet. Yellow: lipids that were not assigned to the extracellular or intracellular leaflets. Grey outline: dDAT. **f)** Performance of the spectral-clustering leaflet detection algorithm on the neuronal membrane. Cyan: extracellular leaflet. Dark blue: intracellular leaflet. Grey outline: dDAT.
